# Plpp3 regulates craniofacial morphogenesis but is dispensable for establishing the midbrain-hindbrain boundary

**DOI:** 10.64898/2026.10.09.757845

**Authors:** Arlen Ramírez-Corona, Elliot Flores-Oliva, Sheyla Cruz-Cruz, Ana Valeria Martinez-Silva, Jocelyn Vázquez-Díaz, Angélica Piedra-Ramírez, Ana Sofía Chávez-Ramos, Erick A. Rojas-Pacheco, Diana Escalante-Alcalde

## Abstract

Phospholipid Phosphatase 3 (PLPP3), an integral membrane enzyme, is essential for embryonic development. It plays a critical role in lipid metabolism and in regulating the cell signaling of several bioactive lipid mediators. One of these lipids, LPA, has been suggested to participate in establishing the midbrain-hindbrain boundary (MHB) and in cranial neural tube closure in mice, and in neural crest migration in frogs. To uncover PLPP3’s role in these processes, we studied the effects of global and conditional *Plpp3* inactivation in *Wnt1::Cre2*-expressing domains. We show that *Plpp3* expression is dispensable for establishing the MHB, although our data support the involvement of LPA-mediated signaling in regulating *Fgf8* expression during anterior embryo development. Its deficiency in neural crest cells yields craniofacial dysmorphology and neonatal lethality due to cardiac failure. Craniofacial defects include a reduced mid frontonasal process, hypotelorism, and an anterior shift of the coronal suture position, driven by localized cell death between E10.5 and E11.5. Our work reveals novel functions of PLPP3 in regulating craniofacial development and coronal suture positioning.

**Highlights:**

- PLPP3 is dispensable in the whole embryo for establishing the midbrain-hindbrain boundary, but it regulates cephalic *Fgf8* expression levels.
- Loss of *Plpp3* in pre-migratory neural crest cells causes frontonasal hypoplasia, hypotelorism, and abnormal coronal suture positioning.
- Localized cell death around E10.5 precedes craniofacial defects.

## Introduction

<u>P</u>hospholipid <u>P</u>hos<u>p</u>hatases (PLPPs, also known as PPAP2s or LPPs) are cell-surface and intracellular membrane-bound integral proteins with six transmembrane domains (Kok et al., 2012). They form homo- and hetero-tetramers (Wang et al., 2026; Yang et al., 2026) with channel-like structure that hydrolyze bioactive lipids derived from phospholipids and sphingolipids, such as lysophosphatidic acid (LPA), sphingosine 1-phosphate (S1P), phosphatidic acid (PA), and ceramide 1-phosphate (C1P) (Kok et al., 2012).

Among the three mammalian PLPPs that dephosphorylate a wide range of lipids, PLPP3 is unique since it is the only one essential for embryonic development, regulating extra-embryonic vascularization, placenta formation, axis patterning, cardiovascular development, and endothelial barrier integrity (Chatterjee et al., 2016; Escalante-Alcalde et al., 2003; Panchatcharam et al., 2014). Furthermore, it possesses an Arg-Gly-Asp (RGD) motif in humans and an Arg-Gly-Glu (RGE) motif in mice in the second extracellular loop that enables interaction with αvβ3 and α5β1 integrins, directly promoting cell-cell adhesion and integrin-mediated signaling (Humtsoe et al., 2005; Humtsoe et al., 2003). This feature, independent of its catalytic activity, is important for endothelial cells although the relevance of this motif has not been established *in vivo* yet.

As cell-surface proteins, PLPPs regulates extracellular LPA and S1P levels, therefore receptor-mediated signaling (Kok et al., 2012). PLPP3 has been demonstrated to shape fine concentration gradients of these lipids (Ramos-Perez et al., 2015; Susanto et al., 2017). Additionally, PLPP3 located to the plasma membrane participates in exogenous S1P cellular uptake and recycling into the sphingolipid synthesis pathway (Kono et al., 2022). All PLPPs can also dephosphorylate intracellular substrates either formed downstream of extracellular signaling activation or in organelles such as Golgi and endoplasmatic reticulum (ER). PLPP3 deficient primary mouse embryo fibroblast showed increased extracellular LPA and intracellular PA and reduced intracellular DAG (Escalante-Alcalde et al., 2003), although each enzyme seems to work in a context dependent manner. For instance, overexpression of PLPP3 in HEK293 cells reduces intracellular S1P (Long et al., 2005) while PLPP3 in the ER prevents the accumulation of PA in the mitochondria further impacting fission and mitochondrial bioenergetics (Luo et al., 2024; Yu et al., 2026).

Dysregulation of PLPP3 has been associated with several diseases. A SNP in an intronic regulatory element of *PLPP3* was first associated with increased risk of developing coronary artery disease (CAD) (Krause et al., 2018; Schunkert et al., 2011). By now, genome-wide association studies (GWAS) have reported 208 associations exhibiting significant pleiotropy across several broad trait categories including myocardial infarction, angina pectoris, systolic pressure, decrease HDL cholesterol, blood cell counts, bone mineral density, inflammatory biomarkers, amongst others (https://www.ebi.ac.uk/gwas/home; (Cerezo et al., 2025). Animal models have shown that PLPP3 expression protects against vascular disease by regulating vascular permeability, inhibiting inflammation and limiting intimal hyperplasia after vessel injury (Mueller et al., 2019; Panchatcharam et al., 2013; Panchatcharam et al., 2014). PLPP3 had also been studied in the context of cancer, where its downregulation is mostly associated with enhanced LPA- and S1P-mediated signaling (Tang and Brindley, 2020). The wide spectrum of cellular processes associated to PLPP3 function, makes imperative to understand how it is that it participates in regulating the different processes in which it is involved.

In this work we aimed to analyze if *Plpp3* expressed in the prospective midbrain hindbrain during early mouse embryo development (Escalante-Alcalde et al., 2009) contributed to midbrain-hindbrain boundary organizer (MBH) development, since extracellular LPA-synthesis by *Enpp2*/Autotaxin was suggested to be required for establishing the MHB in the mouse embryo (Koike et al., 2011). Our data supported that indeed LPA-mediated signaling contributes to the establishment of the MHB in *Enpp2^-/-^*embryos and suggested that PLPP3 could play a role in neural crest development. To approach this question we conditionally inactivated *Plpp3* in pre-migratory neural crest cells using the *Wnt1::Cre2* driver. Its deficiency in neural crest-derivatives yields craniofacial dysmorphology and neonatal lethality due to cardiac failure. Craniofacial defects are characterized by anterior shift in the coronal suture position and a mild facial microform of holoprosencephaly. These phenotypes are associated to increased cell death between E10.5-11.5 mainly in regions where NC/mesoderm interacts. Our work unveils novel functions of PLPP3 in regulating craniofacial development and positioning of the coronal suture, and opens new avenues to understand the role of PLPP3 in mammalian development and in human diseases.

## Results

### PLPP3 is not required for establishing the MHB

For this part of the study, we analysed embryos with global inactivation of *Plpp3* using a *lacZ*-reporter allele previously described (Escalante-Alcalde et al., 2009) backcrossed into the CD1 strain to reduce the early gastrulation phenotype, which was less penetrant in this background. Also, given that the intensity and extension of expression domains of MHB genes varies with the developmental stage, all comparisons of markers were performed in stage matched 8-somite embryos.

First, we analysed the expression of the transcription factor *Otx2,* which participates in the patterning of the forebrain and midbrain and in positioning the MHB (Matsuo et al., 1995; Wurst and Bally-Cuif, 2001). *Otx2* expression in *Plpp3^+/+^* (n=12) embryos was as expected in the anterior neural folds, from the prospective forebrain up to the caudal limit of the midbrain, where a sharp border abuts the prospective hindbrain (Fig. 1A). Likewise, although morphological defects in the anterior neural folds of *Plpp3^lacZ/lacZ^* embryos were observed (n=11), a normal *Otx2* expression pattern was present, suggesting a proper induction and placement of the MHB and midbrain-hindbrain patterning. Induction of the MHB is characterized by the expression of *Pax2/5/8*, *En1/2* transcription factors and the morphogens *Wnt1* and *Fgf8.* With this in mind, we next evaluated the expression of the pair domain transcription factor *Pax2*, which is the earliest gene expressed in the prospective MH regions and is later restricted to a domain spanning the posterior midbrain and anterior hindbrain; its expression precedes other regulatory genes important for the induction and maintenance of the MHB organizer activity (Rowitch and McMahon, 1995). Since *Pax2* is also expressed in anterior neural folds, otic placode and pronephric anlagen, we also analysed expression in these structures as a control of the tissue condition. In contrast to the expected pattern found in *Plpp3^+/+^* embryos (n=18, Fig. 1B), variations in *Pax2* expression in the cephalic region of *Plpp3^lacZ/lacZ^* embryos were observed (n=10). In the developing eye, intensity of *Pax2* staining was diminished, while in the prospective MHB, intensity and distribution of signal was variable although present, being weaker and less defined near the neural fold margins (Fig. 1B). The latter suggested that absence of *Plpp3* expression in the 8-somite embryo supports enough *Pax2* expression to initiate the formation of the prospective MHB. *Pax2* expression in the otic placode was variable between the evaluated embryos, a few had normal pattern and expression levels (2/10), while in others *Pax2 in situ* signal seemed compacted and fainter (5/10) or completely absent (3/10) (Fig. 1B). Expression in the pronephric anlagen showed no significant variations compared to control embryos, suggesting that variations on *Pax2* expression in PLPP3-deficient embryos might be region specific.

**Figure 1.**
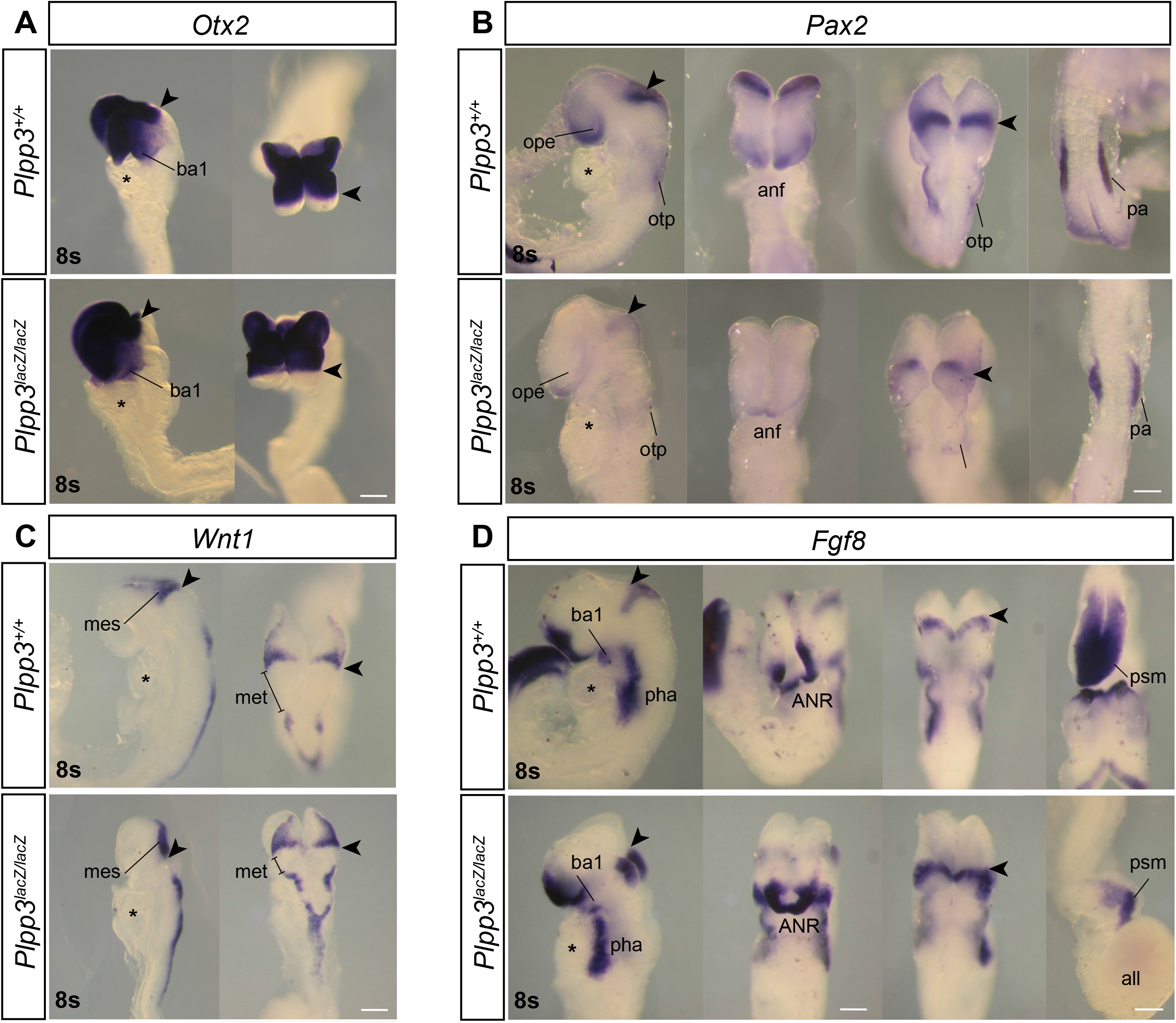
Expression of MHB markers in 8-somite wild-type and *Plpp3^lacZ/lacZ^* embryos. (**A**) *Otx2* expression in lateral (left) and dorsal (right) views. Arrowheads show the position of the MHB. *Otx2* is expressed in the forebrain and the midbrain with its caudal limit before the anterior hindbrain at the MHB. Expression is also observed in the first branchial arch (ba1). *Plpp3^+/+^* (n=12) and *Plpp3^lacZ/lacZ^* (n=11). *, heart. Scale bar 200µ. (**B**) *Pax2* expression decreases in the ANR and MHB in *Plpp3^lacZ/lacZ^* embryos. Images show, from left to right, lateral, frontal and dorsal views of the *Pax2* expression domains. Expression is observed in the neural folds of the future forebrain (*), the MHB region (arrowhead), the future otic vesicle and pronephric anlagen (far right). *Plpp3^+/+^* (n=18) and *Plpp3^lacZ/lacZ^* (n=10). *, heart; otp: otic placode; ope, optic eminence. (**C**) *Wnt1* expression domains appeared mildly extended in *Plpp3^lacZ/lacZ^* embryos. Images show lateral (left) and dorsal (right) views of *Wnt1* expression domains. The anterior end of the embryo is on top. Arrows show the position of the MHB. *Wnt1* is expressed in the caudal limit of the midbrain, along the dorsal margin of the neural folds and on the dorsal midline of the neural tube. No expression is observed at rhombomere 1 (r1). *Plpp3^+/+^* (n=27) and *Plpp3^lacZ/lacZ^* (n=20). *; heart. met; metencephalon. (**D**) *Fgf8* expression in *Plpp3^lacZ/lacZ^* embryos is significantly increased in the AER and MHB but reduced at the caudal domain. Images show, from left to right, lateral, ventral, dorsal and caudal views of *Fgf8* expression domains. The anterior end of the embryo is on top. Lateral view shows *Fgf8* expression domains in the MHB (arrowhead), the first branchial arch (ba1) and pharyngeal tissue (pha). Ventral view shows the extension of the *Fgf8* expression domain along the anterior neural ridge of the forebrain (ANR). Dorsal views show the position of the MHB and the expression of *Fgf8* on the anterior boundary of the hindbrain. Caudal expression of *Fgf8* in *Plpp3^lacZ/lacZ^* embryos showing abnormal presomitic mesoderm (psm) formation. *Plpp3^+/+^* (n=23) and *Plpp3^lacZ/lacZ^* (n=19). all; allantois. *; heart. Scale bar = 200 µm.

We then analyzed expression of *Wnt1*, which encodes the first morphogen produced in the prospective caudal mesencephalon and then is restricted to the anterior MHB. In *Plpp3^+/+^* (n=27) embryos, *Wnt1* expression was on the dorsal neural ridges and the caudalmost domain of the prospective midbrain, neural ridges of the prospective hindbrain (except for the domain corresponding to the metencephalon) and on the roof plate of the closed neural tube (Fig. 1C). In contrast, in *Plpp3^lacZ/lacZ^* embryos (n=20) the intensity of *Wnt1* staining appeared remarkably stronger in the prospective MHB in some embryos (6/20) and more intense or wider on the midbrain (12/20) and rhombencephalon domains (Fig. 1C).

Next, we analysed *Fgf8*, which plays a key role in the patterning activity of the MHB and is restricted to the rostralmost part of the prospective hindbrain. In *Plpp3^+/+^*embryos (n=23), *Fgf8* expression was observed in the anterior neural ridge (ANR), prospective MHB and anterior hindbrain, first branchial arch, pharyngeal region and tailbud (Fig. 1D). In contrast, although the same expression domains were present, the intensity of the staining was evidently stronger in the ANR as well as in the MHB of *Plpp3^lacZ/lacZ^* embryos (n=19). In the latter, the expression domain appeared also wider compared to *Plpp3^+/+^* embryos (Fig. 1D).

To deepen into our hybridization findings, we performed RT-qPCR experiments from mRNA extracted from pools of embryos’ cephalic regions spanning from the developing forebrain to hindbrain, but excluding the heart. As shown in figure 2A, *Fgf8* mRNA was consistently increased up to 5-fold in *Plpp3^lacZ/lacZ^* vs controls confirming that global PLPP3 deficiency causes an increase in cephalic *Fgf8* expression. *Pax2* expression was reduced in *Plpp3^lacZ/lacZ^*, likely reflecting the expression reduction observed in mutant embryos by *in situ* hybridization. *Wnt1* expression was variable between both pools analysed, however the differences were not as dramatic as for *Fgf8*. Scanning electron microscopy (SEM) analyses of mutant embryos showed that cells at the neuroepithelial border, presumably coinciding with *Wnt1*-expressing pre-migratory neural crest cells (NCC), were loosely attached and showed extensive filopodia, suggesting that the expansion of *Wnt1* expression domains observed in neural plate borders of mutant embryos could be due to an abnormal distribution of *Wnt1*-expressing pre-migratory NCC (Fig. 2B, C). The latter pointed to a potential role for PLPP3 in NCC development (see below).

**Figure 2.**
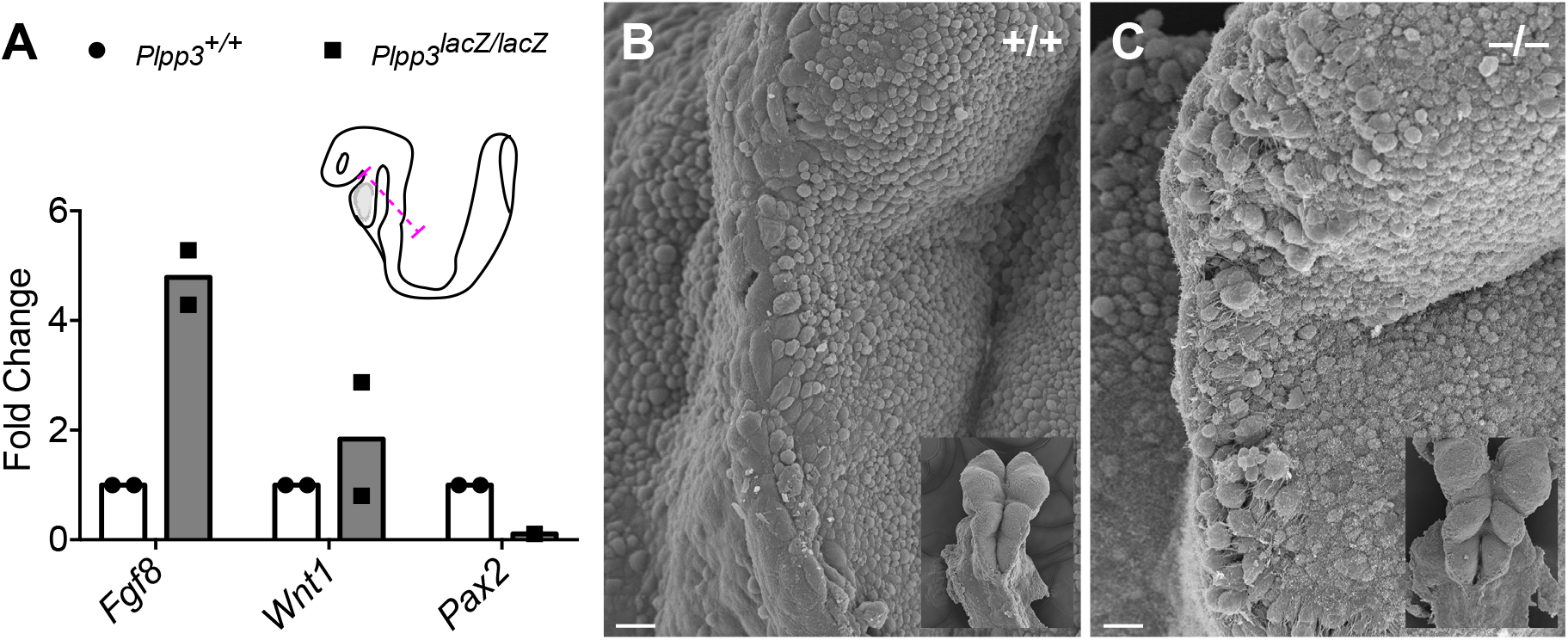
*Fgf8* transcription increases 5-fold *in Plpp3^lacZ/lacZ^* embryos. (**A**) qPCR from 8-somite cephalic regions. The purple line shows the level at which the heads were sectioned. (n=2 independent pools, except for *Pax2*). (**B**) Scanning electron micrographs of the neural plate borders of wild-type and *Plpp3^lacZ/lacZ^*embryos. Note the disorganization of border cells in mutant embryos. Scale bar = 10 µm

Finally, we evaluated the transcription factor *En2,* which is expressed in a region spanning the posterior midbrain and anterior hindbrain. Its expression is regulated by WNT1 and FGF8 and together with *En1* directs midbrain and cerebellum development. *Plpp3^+/+^* (n=12) embryos showed *En2* expression at the MH area, covering the caudal midbrain and the anterior hindbrain (Fig. 3A). *Plpp3^lacZ/lacZ^* embryos (n=11) showed a similar *En2* expression pattern (Fig. 3A). Altogether, our data demonstrate that lack of PLPP3 in the whole 8-somite mouse embryo does not prevent the establishment of the MHB.

**Figure 3.**
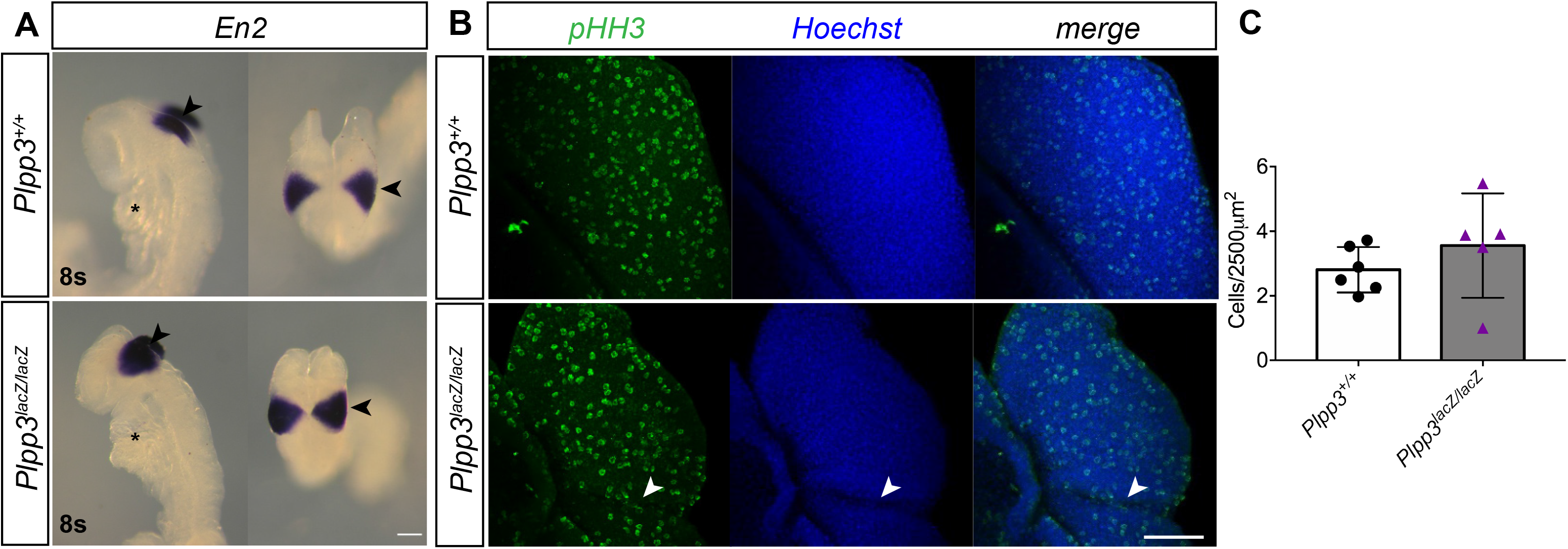
Neither *En2* expression nor proliferation in the neuroepithelium are affected by global *Plpp3* deficiency. (**A**) Images show lateral and dorsal views of the *En2* expression domain. The anterior end of the embryo is on top. *En2* expression is observed in the neural folds of the MHB region, spanning from the posterior midbrain to the metencephalon in the anterior hindbrain. Arrows show the position of the MHB. *Plpp3^+/+^* (n=12) and *Plpp3^lacZ/lacZ^*(n=11). *; heart. Scale bar = 200 μm. (**B**) Stereotypical groves in *Plpp3^lacZ/lacZ^* embryos around 8s don’t seem to be caused by decreased cell proliferation in the MHB region. Images show z-projections of the dorsal views of wild-type and *Plpp3^lacZ/lacZ^*embryos evaluated for pHH3 positive nuclei (green). Nuclei are shown in blue using Hoescht. White arrowheads indicate the position of stereotypical groves. Scale bar = 100 µm. (**C**) Quantification of pHH3 positive cells in the cephalic region of embryos. Error bars show standard error. *= P< 0.0240; **= P< 0.0096. *Plpp3^+/+^* (n=12), and *Plpp3^lacZ/lacZ^* (n=6).

### Cell death patterns in mutant embryos suggest enzyme involvement in NCC development

Consistently, *En2* anterior and posterior expression limits were flanked by stereotypic grooves. To assess the cellular mechanism originating these morphological abnormalities we analysed proliferation and cell death of the neuroepithelium around the MHB area. Cell proliferation was evaluated by wholemount immunofluorescence of phospho-Histone H3 (pHH3). *Plpp3^+/+^* embryos showed a smooth neuroepithelium with uniform distribution of pHH3 positive cells along the MHB area (Fig. 3B). Likewise, pHH3 positive cells in the neuroepithelium of *Plpp3^lacZ/lacZ^* embryos were homogeneously distributed, including the grooves and folds (Fig. 3B). Quantitative analysis of the amount of pHH3 positive cells per 2500 µm^2^ of neuroepithelium revealed no statistically significant difference in proliferating cells in mutant vs wild-type embryos (Fig. 3C). We then evaluated cell death in whole embryos through the detection of active cleaved Caspase-3 (aCas-3). Discrete domains of focalized cell death were observed in *Plpp3^+/+^*embryos (n=7), coinciding with those previously described during normal development (ANR and preotic sulcus of the rhombencephalon; Massa et al., 2009) (Fig. 4A-C). In contrast, the quantity of aCas-3 positive cells observed in the ANR of *Plpp3^lacZ/lacZ^* embryos (n=15, Fig. 4D-F) was strongly reduced when compared with controls of equivalent developmental stages (compare Fig. 4C with 4D). Although the presence of aCas-3 positive cells was observed at the grooves in *Plpp3^lacZ/lacZ^* embryos (Fig. 4E), the number of dead cells does not account for the dramatic changes in morphology that already appear in less developed embryos (Fig. 4D). In more advanced embryos (Fig. 4F), we identified several areas with increased cell death such as the lateral aspects of the anterior cephalic region, the optic sulcus area, the most anterior border and central region of the developing first branchial arch, and in the area just anterior to the otic placode (Figs. 4E and 4F). These regions spatially overlap with known NCC migratory pathways namely, the first branchial arch stream originating from dorsal rhombomeres 1 and 2 and the second branchial arch stream in rhombomere 4. Interestingly, although at E8.25-8.5 NCC do express *Plpp3* at low levels (Fig. S1, Pijuan-Sala et al., 2019), its expression is notably more abundant in rhombomere 3 and the otic placode in rhombomere 5 (Figure 4F, inset). The latter result suggested the tempting idea that PLPP3 could contribute to somehow restricting the pathways of neural crest cells migration, similar to the role of *wunen* in *D. melanogaster* primordial germ cell migration (Starz-Gaiano et al., 2001). Explant migration assays of neural plate borders of 8-somite *Plpp3^lacZ/lacZ^*embryos showed that SOX10^+^ NCC have defective migration (Fig. S2). Our SEM analysis, in combination with altered cell death patterns and abnormal NCC migration in mutant explants, suggested that PLPP3 could play a role in NCC development (see below).

**Figure 4.**
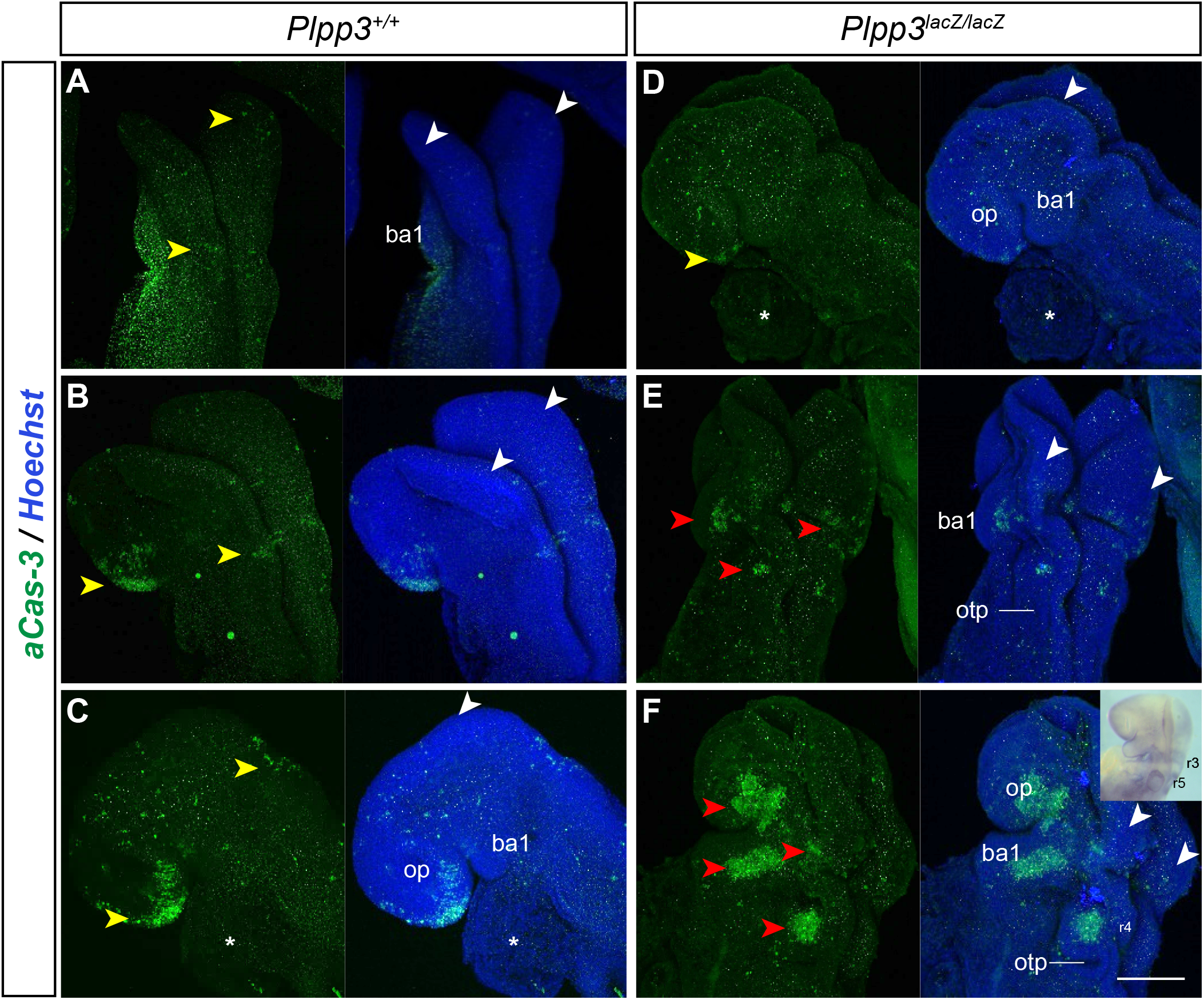
Stereotypical grooves in MHB of *Plpp3^lacZ/lacZ^* embryos are not caused by increased cell death. Lateral and dorsal views of *Plpp3^+/+^* (**A, B, C**) and *Plpp3^lacZ/lacZ^* embryos (**D, E, F**) at different stages of development. Immunofluorescence shows dying cells positive for cleaved caspase-3 (aCas-3 green). White arrowheads show the approximate location of the MHB. Yellow arrowheads show normal location of cell death domains in the caudal rhombencephalon and ANR. Note the reduction of Cas-3 in the ANR of mutant embryos (compare **B** and **D**). Red arrowheads show broader or ectopic cell death domains in the neuroepithelium and lateral cephalic regions of more developed *Plpp3^lacZ/lacZ^* embryos (**F**). Nuclei are shown in blue using Hoechst. *Plpp3^+/+^* (n=7) and *Plpp3^lacZ/lacZ^*(n=15). *; heart. ba1: branchial arch 1; op: optic placode; otp: otic placode; r3: rhombomere 3; r5: rhombomere 5. Inset, *in situ* hybridization of *Plpp3*. Scale bar= 200 µm.

### Absence of local expression of Plpp3 does not alter Fgf8 expression in the MHB

Since the severity of malformations found in PLPP3-deficient embryos beyond 8-somites prevents the accurate analysis of phenotypes in later stages, and since we cannot rule out that some of the alterations described above were due to the aberrant accumulation of a PLPP3 substrate(s) in the amniotic cavity, we sought to conditionally inactivate *Plpp3*. We used the *Wnt1::Cre2* transgenic line (Lewis et al., 2013) to drive recombination of our *Plpp3* conditional allele (*Plpp3^f/f^*; Escalante-Alcalde et al., 2007) in the midbrain, dorsal diencephalon, anterior hindbrain as well as in NCC-derivatives (<u>https://</u> www.informatics.jax.org/recombinase/specificity?id=MGI:5485027&system=nervous%20system). We initially obtained embryos from mating *Wnt1::Cre2*; *Plpp3^f/+^*males to *Plpp3^f/f^* females (Figure 5A). In this particular *Cre*-line, the transgene is active in the male germ line (Dinsmore et al., 2022). Due to this characteristic, embryos that are homozygous mutants in the structures of interest are heterozygous in the rest of their cells (*Wnt1::Cre2*;*Plpp3^f/Δf^*). This feature allowed us to assess if the local expression of *Plpp3* is required to regulate *Fgf8* expression, taking into account that expression of *Wnt1* and *Fgf8* partially overlap during the emergence of the organizer (Canning et al., 2007), that this Cre-line is transiently active in a broader region that includes the anterior hindbrain/cerebellum and that *Fgf8* expression in the MHB of *Plpp3^lacZ/+^* embryos was not modified (not shown).

**Figure 5.**
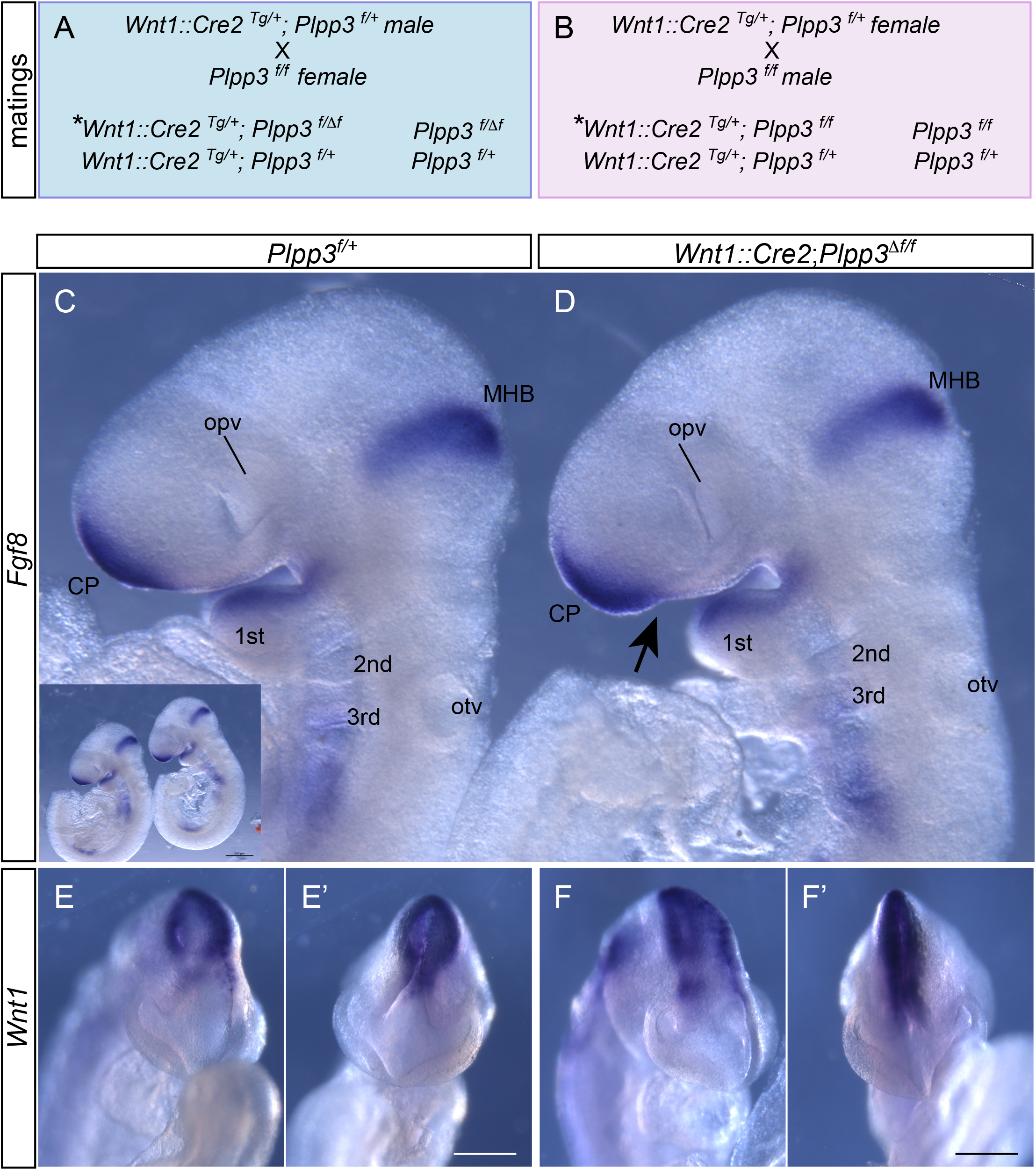
Conditional inactivation of *Plpp3* via males carrying the *Wnt1-cre2* transgene, does not yield *Fgf8* overexpression in the MHB or commissural plate. (**A**) Resulting genotypes from males carrying the *Wnt1::Cre2* transgene. Note that males inherit alleles already recombined (ΔF), therefore with heterozygous expression of *Plpp3* in the whole embryo. (**B**) Resulting genotypes from females carrying the *Wnt1::Cre2* transgene. Under this condition *Plpp3* is only inactivated in *Wnt1*-expression domains. *Fgf8 in situ* hybridization of control (**C**, *Plpp3^f/+^*) and mutant (**D**, *Wnt1::Cre2;Plpp3^ΔF/F^*) E9.5 embryos produced from mating males carrying the transgene. No change in *Fgf8* expression was observed in mutant embryos, although a small depression (black arrow) was observed ventral to the *Fgf8* anterior expression domain. *Wnt1* in situ hybridization of control (**E**, **E’**) and mutant (**F**, **F’**) E9.5 embryos produced from mating males carrying the transgene. ¾ (**E**, **F**) and frontal (**E’**, **F’**) views. *, mutant embryos from each mating scheme; 1^st^ 2^nd^, 3^rd^ brachial arches; cp, commissural plate; MHB, midbrain/hindbrain boundary organizer; opv, optic vesicle; otv, otic vesicle. *Fgf8* inset scale bar= 500 μm; *Wnt1* scale bar= 200 μm.

We recovered twenty E9.5 embryos out of 3 pregnant females. All embryos appeared normal and with complete neural tube closure. After genotyping using yolk sac DNA, six were *Plpp3^f/+^* (*f/+*, 30%), five *Wnt1::Cre2*;*Plpp3^f/+^* (*Cre;f/+*, 25%), five *Plpp3^f/Δf^* (*f/Δf*, 25%) and four *Wnt1::Cre2*;*Plpp3^f/Δf^* (*Cre;f/Δf*, 20%), showing normal Mendelian distribution of genotypes and survival of mutants at this stage (*Cre;f/Δf*). Embryos were subjected to *Fgf8 in situ* hybridization and expression patterns compared between genotypes. No evident difference in *Fgf8* expression in the commissural plate (CP) or MHB was observed between *Cre;f/Δf* and *Cre;f/+* (not shown) or *f/+* controls (Fig. 5C, D). Also, no difference in *Wnt1* expression was detected between genotypes (Fig. 5E, F). These data show that inactivation of *Plpp3* with the *Wnt1::Cre2* driver inherited via males does not affect *Fgf8* expression in the MHB or CP.

### Ablation of Plpp3 with the Wnt1::Cre2 driver produces craniofacial dysmorphology

To further investigate the role of PLPP3 during cephalic development, and to rule out a possible additive effect of whole embryo heterozygous ablation of *Plpp3* in *Cre;f/Δf* individuals (Fig. 5A), we mated *Wnt1::Cre2*; *Plpp3^f/+^*females to *Plpp3^f/f^* males (Fig. 5B). We noticed that about one-quarter of the littermates died soon after birth (P0), some within the next 24 to 48 hours, and very few *Wnt1::Cre2*; *Plpp3^f/f^*individuals reached or survived beyond weaning, depending on the genetic background (Tables 1 and 2). Phenotypic analysis of P0 pups revealed that a portion of the individuals lacked milk spot, were cyanotic, showed gasping or irregular breathing, a sharper snout and more noticeable eyes. Genotyping showed that those individuals corresponded to *Wnt1::Cre2*; *Plpp3^f/f^* mutants (CKO) and were present in expected Mendelian ratio (Tables1 and 2). Their palate was properly fused, however their secondary palate appeared mildly arched with rugae abnormally oriented (not shown). No suckling defects were detected suggesting that cardiac failure, instead of a feeding impairment, was the main cause of lethality. Characterization of the cardiovascular phenotype will be described elsewhere (Rojas-Pacheco *et al*., in preparation).

**Table 1.**
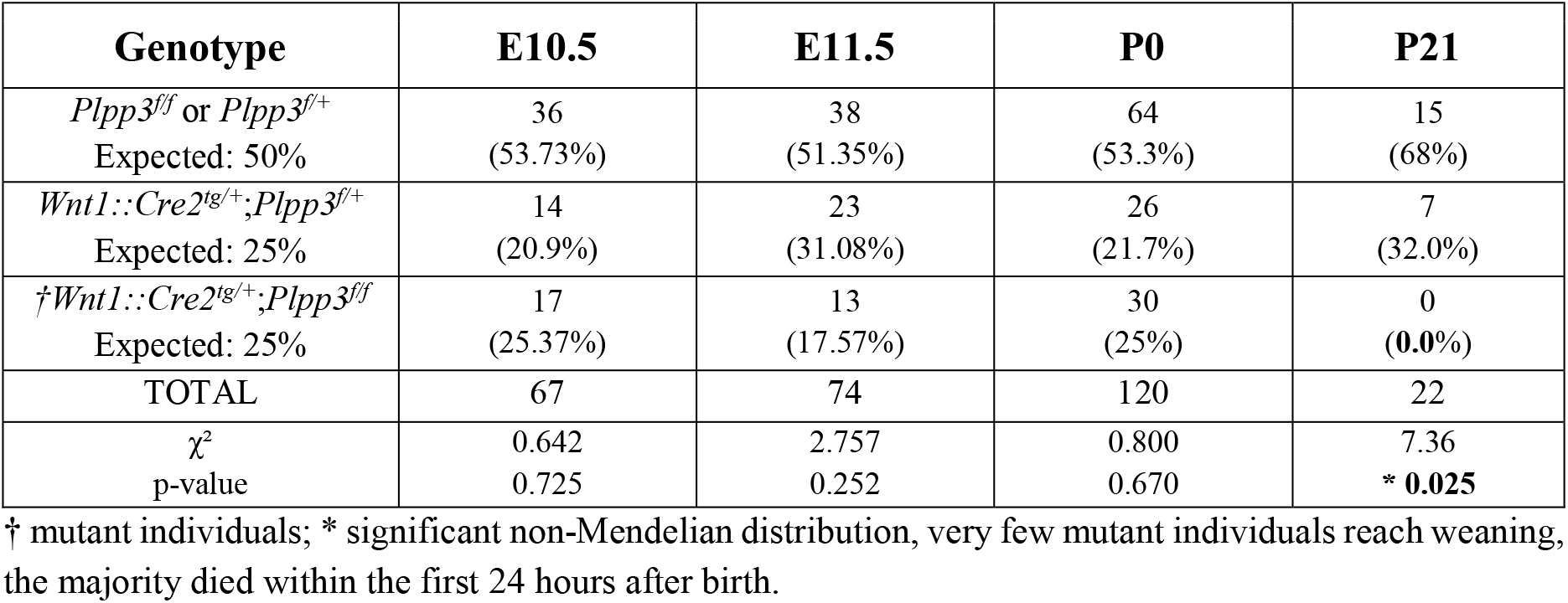
Distribution of genotypes with Cre inherited via females (C57BL6/J).

**Table 2.**
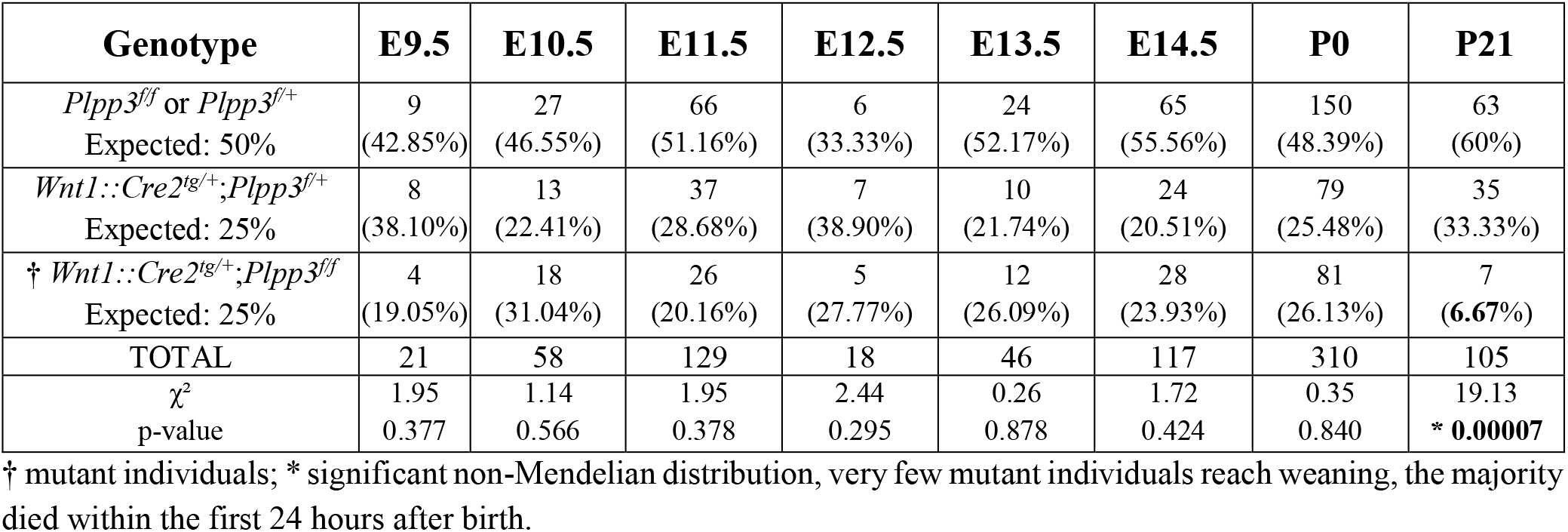
Distribution of genotypes with Cre inherited via females (C57.129).

To study the consequences of *Plpp3* inactivation on the skull structure, P0 heads from control and CKO individuals were stained with alcian blue and alizarin red. *Wnt1::Cre2*; *Plpp3^f/f^*skulls consistently showed a smaller frontal bone and anterior expansion of the parietal bone with the consequent anterior shift of the coronal suture, and lack of tissue in the most anterior midcentral part of the interparietal bone (Figs. 6A, D; S3 and landmarks table S1). Distances between points shown in figure 6 B and E demonstrated that in CKO, frontal bone length is strongly reduced, and the anterior aspect of the frontal bone was narrower than wildtype or heterozygous controls (Figs. 6G-H and S3). The change in these features produced an anterior face dysmorphism in CKO, making the angle formed by the anterior mid central point of the frontal bone (point F2) and the most lateral points of the coronal suture (points F9, F10) 20° bigger than in control individuals (Fig. 6B, E, I). These morphological changes are responsible for an inward shift in the direction of the eyes. Surprisingly, no significant reduction in the length of the nasal and maxillary bones was observed, despite nasal bones being narrower and the maxillary bones mildly undergrown (Figs. 6 and S3). Analyses of lateral views of crania also showed the evident anterior displacement of the coronal suture, and a mild downward displacement of the parietal-squamosal suture in CKO skulls. Furthermore, small ectopic ossification centres were sometimes observed in the fronto-nasal and parietal-squamosal sutures (Fig 6F, arrows in inset) and occasionally in the interfrontal space.

**Figure 6.**
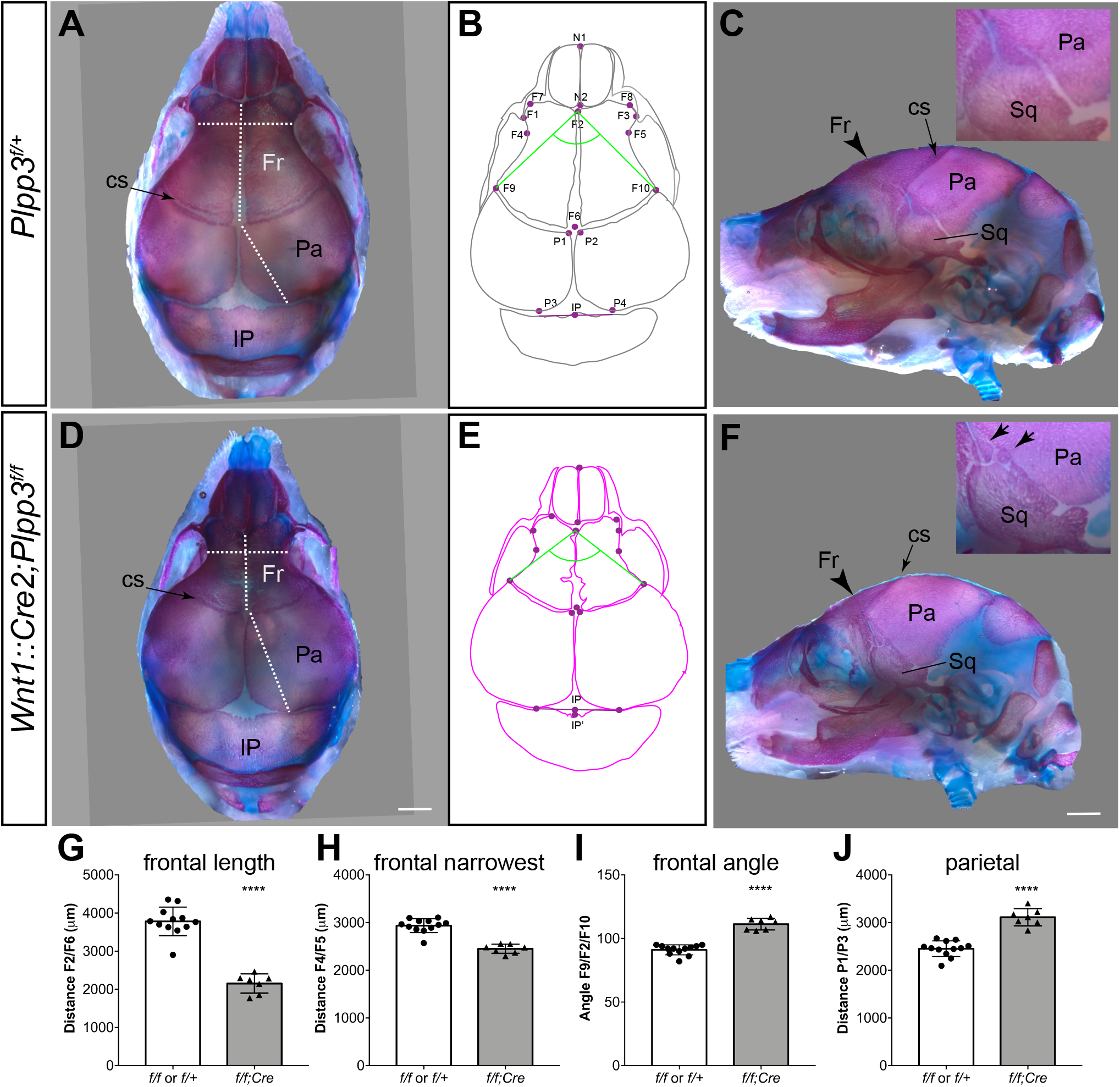
Conditional inactivation of *Plpp3* in NCC, yields anterior dysmorphology and reduction of the frontal bone size in neonates. Skeletal preparations in dorsal (**A**, **D**) and lateral (**C**, **F**) views of control (**A-C**) and CKO (**D-F**). (**B**, **E**) Schematic representation of skulls and landmarks (see also Table S1) used for morphometric analysis. Note the dramatic decrease of the anterior frontal bone length (**G**) and width (**H**) in CKO. This produces an inward change in the position of the eyes represented by the angle formed between. F9/F2/F10 landmarks (**I**, green angle). (**J**) Reduction of the frontal bone length is concomitant with anterior expansion of the parietal bone in CKO. The parietal squamosal suture is mildly displaced downward and small ectopic ossification centres were sometimes observed in the frontal/squamosal/parietal suture (arrows in the inset of **F**). Complete measurements are in Fig. S3, since the 3 functional genotypes were assessed, Dunnett’s one-way ANOVA was applied to the data, although only *f/f* or *f/+* controls and *f/f;Cre* were graphed in this figure ****P>0.0001. cs, coronal suture; Fr, frontal bone; IP, interparietal bone; Pa, parietal bone; Sq, squamosal bone. Scale bar= 1000 µm.

### Plpp3 expression in NCC contributes to positioning of the coronal suture

To investigate the distribution of NCC in relation to the appearance of the craniofacial dysmorphology and abnormal frontal and parietal bone development, we performed lineage tracing experiments using a Cre/double-fluorescent reporter line (Muzumdar et al., 2007). Upon Cre expression, recombined cells express membrane-sorted EGFP (mG), while the unrecombined cells express membrane-sorted tdTomato (mT). Analysis of E14.5 CKO individuals showed reduced distance between the eyes (hypotelorism) and a narrower snout (Fig. 7A, B, G, H and S4); the posterior border of the NC-derived frontal tissue was displaced anteriorly relative to the eye, and midfrontal and frontal NC-derived tissue was hypoplastic (Fig. 7C, D). Also, mesoderm-derived tissue positive for tdTomato was more prominent and extended in the periorbital area, and the boundary between NCC and mesoderm in the prospective squamosal bone area was smoother and mildly displaced downward (Fig. 7C, D). These data indicated that in CKO individuals, interactions between facial NCC and mesoderm apposition fronts might be perturbed and that PLPP3 expression in facial NCC is required for the proper positioning of the coronal suture. Alkaline phosphatase staining of the same embryos corroborated the anterior displacement of the boundary between the developing frontal and parietal osteogenic domains, since the ossification domains corresponding to the developing frontal and parietal bones were reduced and expanded, respectively, and the coronal suture mesenchyme was positioned forward (Fig. 7E, F, I).

**Figure 7.**
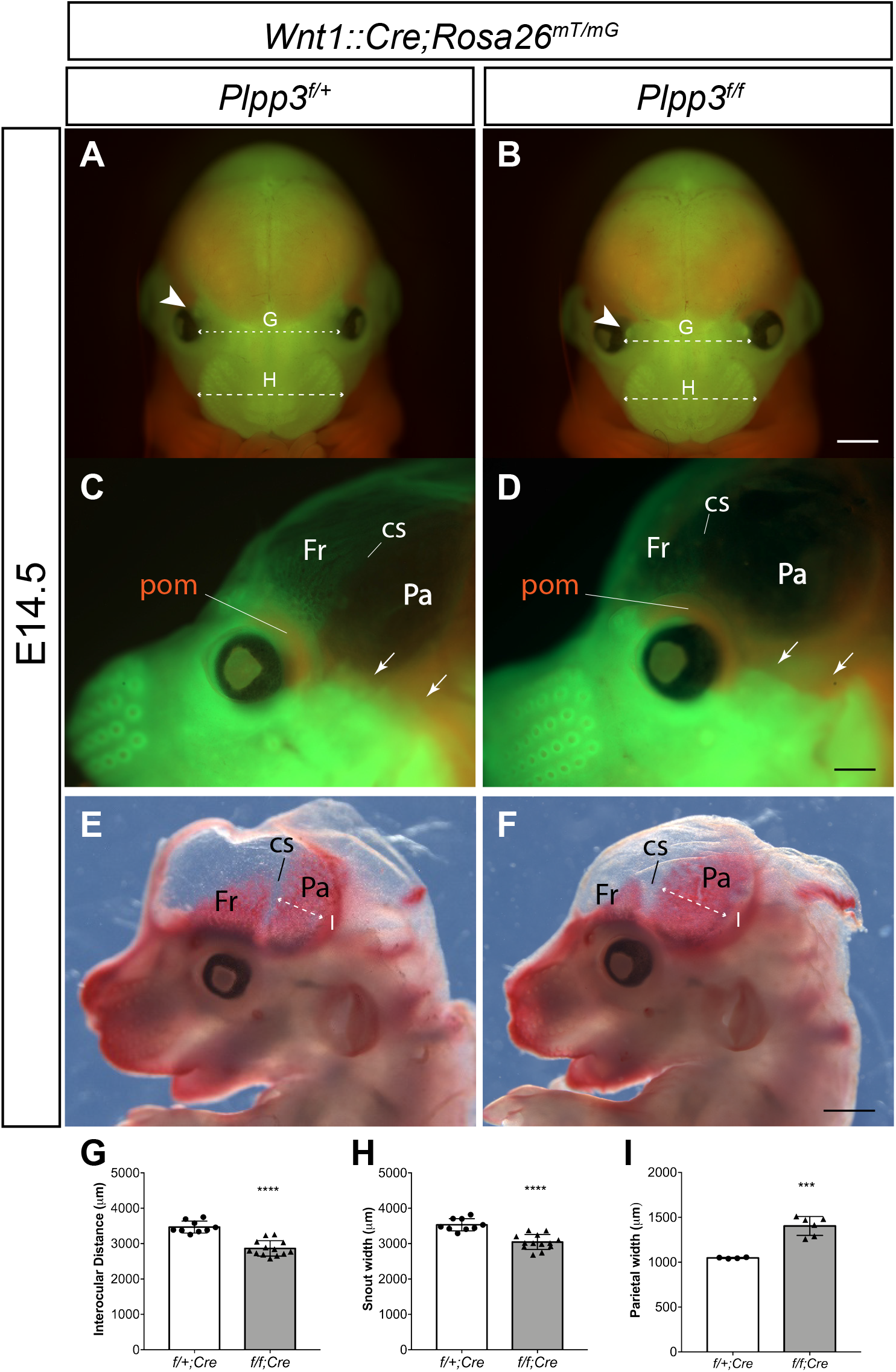
Hypotelorism and anterior shift of the coronal suture position in E14.5 CKO. Lineage tracing in control (**A**, **C**) and CKO (**B**, **D**) E14.5 embryos. EGFP+, *Wnt1::Cre2* mediated recombination; tdTomato+, unrecombined. Note that in the CKO, anterior displacement of supraorbital follicles was observed (white arrowheads) and the distance between the eyes and the snout width are smaller than in controls (**A**, **B**, **G**, **H**). (**C**, **D**). Lateral views of midsagittally-sectioned heads, brain removed, showing reduced EGFP+ frontal tissue with anterior displacement of the coronal suture position and periorbital tdTomato+ mesenchyme of mesodermal origin. Small arrows show lack NC-derived tissue in the temporal area of CKO. (**E**, **F**, **I**) Alkaline phosphatase staining of the sectioned heads corroborates the anterior shift of the coronal suture position and anterior expansion of the parietal domain. cs, coronal suture; Fr, frontal; Pa, parietal; pom, periorbital mesenchyme. (G, H) Student’s *t* test **** P>0.0001, control n=9, CKO n=13. (I) Welch’s *t* test *** P 0.0004, control n=4; CKO n=6. (A, B, E, F) Scale bar= 1000μm; (C, D) Scale bar= 500 μm.

To delve into the mechanism(s) that contributes to the shift in the position of the coronal suture, we initially analysed the number of proliferating and dying cells in CKO and control lineage traced embryos at E14.5 (Fig 8). At this stage, we found no evidence of changes in proliferation or cell death in the neighborhood of the NC/mesoderm boundaries of CKO embryos (Fig. 8). However, two very important findings were made in CKO. At the posterior supraorbital mesenchyme (SOM) level, besides the parietal bone primordium, the tectum transversum of the chondrocranium (also derived from the mesoderm) was also placed forward (Fig 8A, B). Furthermore, there was a consistent absence of NC-derived dermal cells adjacent to the developing temporalis muscle and just above the NC/mesoderm border (Fig. 8C, D). These observations suggested that not only are the dermal cells that will form the frontal bone primordium missing, but also other NC-derived components that border the mesoderm. The latter raised the possibility that in some NC/mesoderm boundaries, *Plpp3*-deficient dermal cells could be dying. To test this hypothesis, we initially traced back the earliest time point where abnormalities were detected. Alterations in head morphology were evident in lateral views of CKO embryos at E11.5, showing a shorter distance between the most distal part of the frontonasal process and the eye. Also, the dorsal borders of NC-derived tissue in the first and second pharyngeal arches appeared smoother, allowing a better observation of the anterior aspect of the trigeminal ganglion (Fig. 9A, B). In frontal views, anterior head and frontonasal processes of CKO appeared narrower than controls and hypotelorism with a more inward position of eyes was already evident (Fig 9C-D, S4). Although reduction in the size of telencephalic vesicles was evident at this stage, no obvious thinning of the neuroepithelium was observed in CKO (Fig. 9C’, D’) suggesting abnormal ventricular expansion due to impaired cerebrospinal fluid accumulation or a physical constriction imposed by abnormal expansion of the NCC population (i. e. meninges).

**Figure 8.**
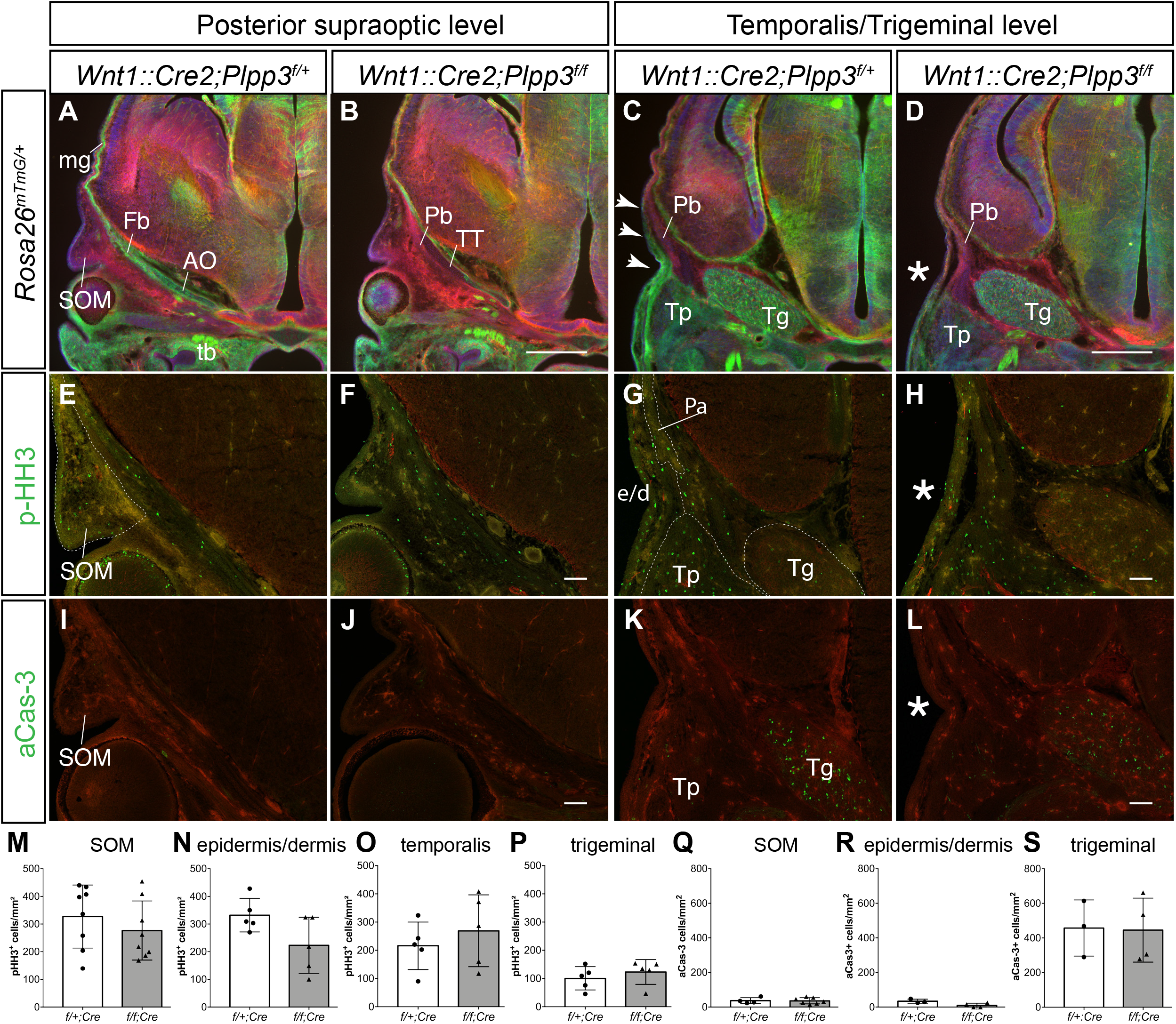
Lack of *Plpp3* expression in NC-derivatives also alters the chondrocranium and temporal dermal cells. All analyses were performed in E14.5 embryos. (**A-D**) EGFP/tdTomato-lineage tracing in coronal sections at the posterior eye (**A**, **B**) and temporalis muscle/trigeminal ganglion (**C**, **D**) levels. (**A**, **B**) Remarkably, in CKO individuals, not only the parietal bone is placed at this eyes level, but the mesoderm-derived tectum transversum of the chondrocranium. (**C**, **D**) Evident loss of dermal EGFP+ tissue was observed in the temporal NC/mesoderm boundary. Compare distribution of tissue in the control (white arrowheads) with equivalent region in the CKO (*). Immunofluorescence against phospho-Histone H3 (**E-H**) and active Caspase-3 (**I-L**) to assess for differences in cell division and cell death, respectively. The analysed areas are indicated by dashed lines. Quantification of marker+-cells in the supraorbital mesenchyme (**M**, **Q**), epidermis/dermis area (**N**, **R**; e/d), temporalis muscle (**O**; Tp) and trigeminal ganglion (**P**, **S**; Tg). aCas-3, active Caspase-3; AO, ala orbitalis; e/d, epidermis/dermis area; Fb, frontal bone; mg, meninges; pHH3, phospho-Histone H3; Pb, parietal bone; SOM, supraorbital mesenchyme; tb, tooth bud; Tg, trigeminal ganglion; Tp, temporalis muscle; TT, tectum transversum. Student’s *t* test. No significant differences were detected. pHH3: SOM n=8, other regions n=5. aCas-3: SOM control n=6, CKO n=7, other regions control n=3, CKO n=4. (A-D) Scale bar=500 µm; (E-L) Scale bar=100 µm.

**Figure 9.**
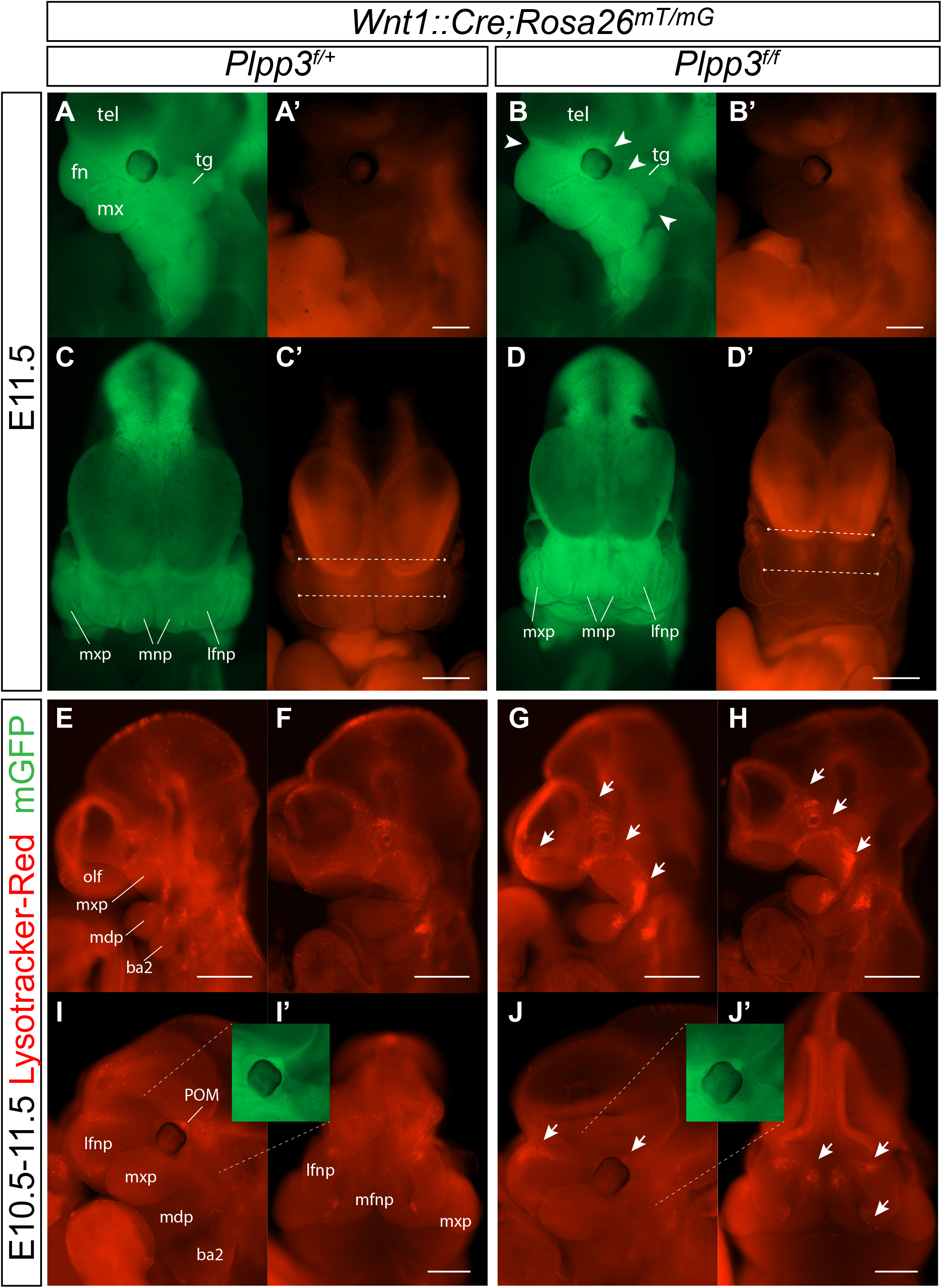
First evident phenotypic features are traced back to E11.5 and preceded by localized increases in cell death. E11.5 control (**A**, **C**) and CKO (**B**, **D**) lineage traced embryos in lateral and frontal views of EGFP (green) and tdTomato (red’) channels. (**A**, **B**) In the CKO, a deeper frontonasal depression, lack of posterior periorbital EGFP+ cells and lack of superficial NC-derived tissue in the dorsal border of branchial arches makes their borders appear smoother (arrowheads). At this stage, a significant reduction in the interocular distance and nasal processes width was observable (see Fig. S4C for measurements). Control (**E**, **F**) and CKO (**G**, **H**) E10.5 embryos stained with LysoTracker. Increased cell death was observed in the frontonasal, supraorbital and suborbital regions as well as in the dorsal and hinge areas of the first branchial arches (white arrows). (**I**, **I’**, **J**, **J’**) Later, at E11.5, augmented cell death was still observed in the proximal lateral and medial nasal prominences, while cell death in the periorbital mesenchyme appeared reduced (white arrows). Inserts: EGFP-channel of the same embryos showing the loss of posterior periorbital mesenchyme. ba2, 2^nd^ branchial arch; fn, frontonasal process; lfnp, lateral frontonasal prominence; mdp, mandibular process; mnp, medial nasal prominence; mxp, maxillary process; olf, olfactory placode; POM, periorbital mesenchyme; tel, telencephalon; tg, trigeminal ganglion. Scale bars= 500 µm.

Then, we stained E10.5-E11.5 lineage traced embryos with LysoTracker Red to probe for abnormal cell death patterns (Fogel et al., 2012). Remarkably, we noticed a strong increase of cell death in some regions of CKO embryos such as the supraoptic area, the hinge area and dorsal border of the 1^st^ branchial arches (E10.5, Fig. 9E-H), the frontal aspect of lateral and medial nasal processes (E10.5-11.5, Fig. 9E-J), all coinciding with structures that will later show altered shape, loss of NC-derivatives or expansion of mesodermal derivatives (Fig. 9A, B, insets in I, J). In some regions, cell death was mildly reduced in E11.5 CKO, such as in the periorbital mesenchymal condensations and the lambdoidal junction, which are also programmed cell death areas required for proper insertion of periocular muscles and primary palate/lip formation, respectively (Fig. 9J, L) (Compagnucci et al., 2021; Sulik et al., 2001).

Our data support the idea that a local increase of cell death in CKO embryos between E10-E11.5 in the supraorbital and temporal areas allows the anterior displacement of mesodermal cells. Between E10.5 and E11.5, *Plpp3* is mildly to strongly expressed in the regions affected by the mutation, and interestingly, *Enpp2, Sphks* and several LPA and S1P receptors are expressed in overlapping or neighbouring but non-overlapping domains (Fig. S5; IMPC, Wilson et al., 2025; Ohuchi et al., 2008; Ohuchi et al., 2007; Yokoyama et al., 2009), raising the possibility that PLPP3 expression in NCC modulates LPA- and/or S1P-mediated signalling for shaping the mouse head and NC/mesoderm boundary interactions.

The region where PLPP3’s gene maps in humans and mice is syntenic (Fig. 10A; Kolishovski et al., 2019) and a recent study characterized the epigenome of human craniofacial development (Wilderman et al., 2018). Visualization of the area containing *PLPP3* shows that it encompasses a craniofacial super-enhancer region active mainly during early craniofacial development (CS13 and CS14 equivalent to E10.5 and E12.5 in mice, respectively). Furthermore, visualization of capture chromatin conformation data shows that *PLPP3* is the single coding gene present in a sub-topologically associated domain containing craniofacial-specific enhancers and super-enhancer regions, with contacts consistent with regulatory interactions involving its promoter (Fig. 10B-D). Altogether, our data demonstrate that PLPP3 in NCC-derivatives is essential for neonatal survival and plays a novel role in regulating mouse craniofacial development and potentially in humans as well.

**Figure 10.**
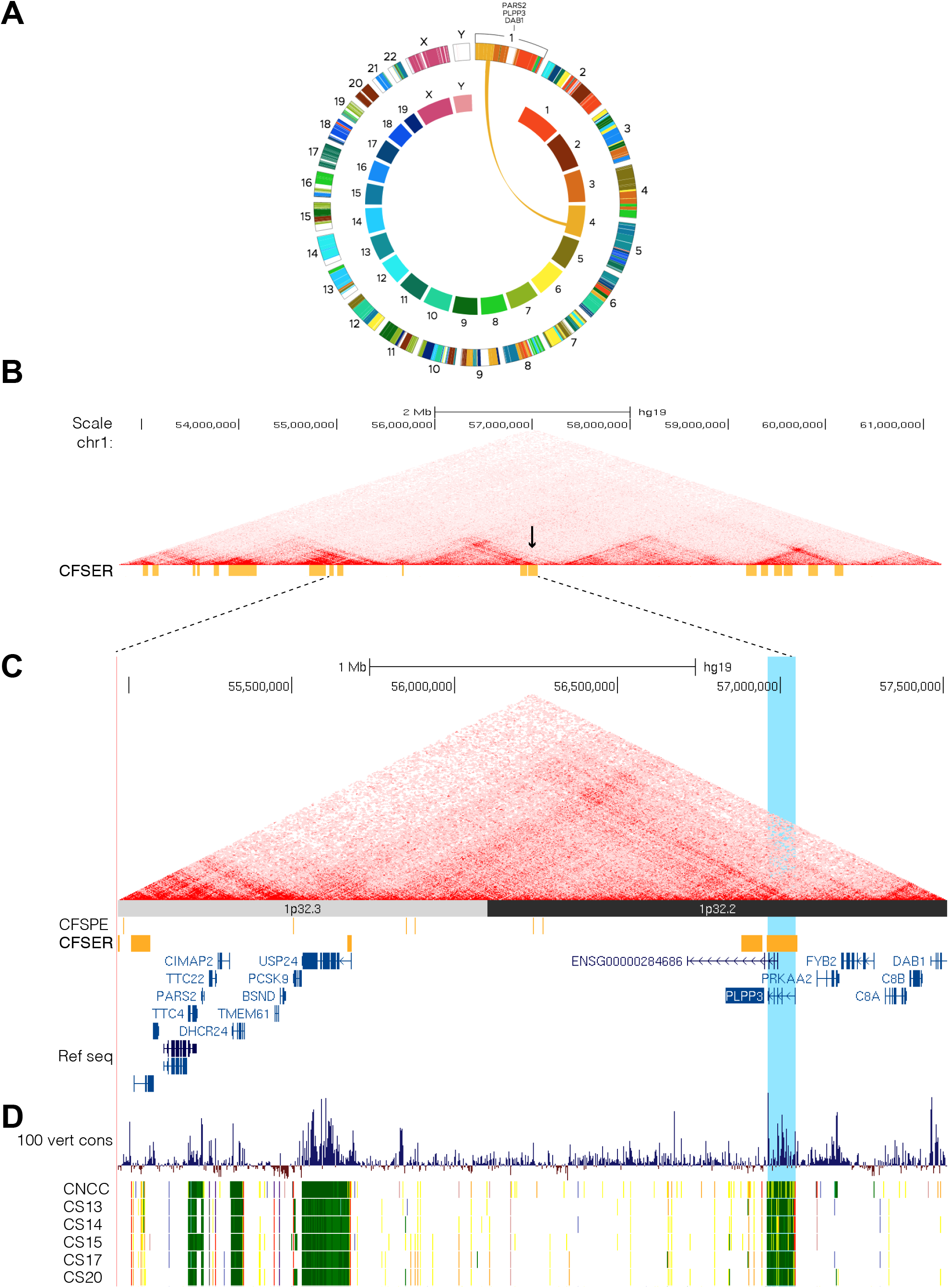
*PLPP3* is the single coding gene within a sub-topological associated domain in humans with active craniofacial super enhancer regions and enhancers. (**A**) Conservation of synteny between mouse and human in the region where *PLPP3* maps. (**B**) Genome browser shot of chromosome 1 showing chromatin architecture in cranial neural crest cells (CNCC) differentiated from hESC, the position of *PLPP3* is indicated by the arrow. (**C**) Zoom-in of 1p32.3 and 1p32.2 regions showing gene and topological organization, position of strong craniofacial specific enhancers (CFSE), strong craniofacial super enhancer regions (CFSER) and vertebrate conservation score. The blue line indicates the position of *PLPP3*. (**D**) Twenty-four state chromatin state segmentations of CNCC and primary human craniofacial tissue from Carnegie stages 13 through 20 (CS13-CS20). *PLPP3* is transcriptionally active in CS13-CS15, equivalent to E10.4-E12.5 in mouse.

## Discussion

### Role of PLPP3 in MHB development

It has been suggested that ENPP2, the main LPA synthesizing enzyme, plays a role in anterior forebrain development and establishment of the MHB of the mouse. In *Enpp2^-/-^*embryos, *Fgf8* is absent or extremely reduced in the ANR and MBH of a portion of the embryos (Koike et al., 2011). In contrast, expression of this morphogen was increased in both regions of 8-somite *Plpp3^lacZ/lacZ^* embryos, confirmed by *in situ* hybridization and RT-qPCR. Given the opposing enzymatic activities of ENPP2 and PLPP3 in the extracellular availability of LPA, our observations support the participation of LPA-mediated signalling in the regulation of *Fgf8* expression in the cephalic region. A functional link between LPA-signalling and *Fgf ’s* expression has been previously described in *Xenopus leavis*. Morpholino-mediated inhibition of *enpp2 or lpar6* produces anterior neural tube development defects and reduction of *fgf8* expression in the ANR and MHB. Moreover, neural tube defects caused by *lpar6* downregulation are more severe when fgf8 signalling is inhibited, suggesting cooperation between both molecules for proper telencephalic development (Geach et al., 2014). Interestingly, *lpar6* inhibition in dorsal anterior blastomeres produces a phenotype (reduced eyes and forehead) that is something similar to the ventralizing phenotype caused by injection of *Plpp3* mRNA in the dorsal anterior blastomeres (reduced eyes and forehead, cyclopia and microcephaly; Escalante-Alcalde et al., 2003), condition in which LPA concentrations should be reduced.

In mouse embryos, previous to the onset of *Fgf8* expression in the anterior region, but posterior to the start of *Wnt1* expression in the MH area (E8.0), *Enpp2*, most of LPA receptors and *Plpp1-3* are expressed in cells of surface and neural ectoderm (*Lpar4 >Lpar6>Lpar2>Lpar1>Lpar3*; Pijuan-Sala et al., 2019). By E8.5, when *Fgf8* expression is already established, *Lpar1/2/4/6* are expressed in the anterior neuroectoderm, including the ANR, with an evident stripe of *Lpar4* in the MHB area (Ohuchi et al., 2008; Pijuan-Sala et al., 2019). Interestingly, at the same stages, *Plpp3* expression in the ANR is low or absent, despite the marked increase of *Fgf8* expression observed in this signalling centre in *Plpp3*-deficient embryos, suggesting a non-local regulation of LPA mediated signaling by PLPP3 in this region. Based on the above, PLPP3 may serve as a rheostat of LPA concentration in the amniotic fluid, thereby modulating signalling in the surface and neural ectoderm. Consistent with this interpretation, conditional inactivation of *Plpp3* in *Wnt1::Cre2* expressing domains, did not produce an alteration of *Fgf8* expression in the MHB or commissural plate of E9.5 embryos (Fig. 5). Altogether these data support that *Fgf8* expression in the ANR and MHB is regulated by LPA and that PLPP3 helps to maintain proper LPA-signalling in these signalling centres.

### PLPP3 in craniofacial development

Our work revealed that PLPP3 is an important regulator of craniofacial development. Its deficiency in NCC*-*derivatives produces an anterior shift in the coronal suture position and a mild facial microform of holoprosencephaly, namely, midfacial hypoplasia and hypotelorism without ventral midline brain malformations.

The coronal suture phenotype seems associated with increased cell death in the supraorbital area between E10.5 and E11.5 (Fig. 9). In the mouse, coronal suture progenitors arise from a cephalic paraxial mesoderm population, responsive to SHH, that migrates to the supraorbital mesenchyme (SOM) and expresses the lineage boundary transcription factor EN1 (Deckelbaum et al., 2012). Part of these cells migrate to form the coronal suture mesenchyme while others remain mixed with supraorbital NCC to form the frontal bone. Although smaller, the frontal bone of our CKO is formed, with the coronal suture mesenchyme correctly separating the parietal bone, indicating adequate migration of the paraxial mesoderm required for coronal suture formation.

*En1* also regulates the positioning of the coronal suture, restricting cell movement between the mesoderm/NCC lineage boundary. *En1* inactivation in NCC produces a caudal shift of the frontal osteogenic domain and the presence of NCC in the coronal suture mesenchyme. Furthermore, *En1* inactivation in *Gli1*-expressing cells (SHH-responsive mesoderm) also produces a caudal shift due to reduction in the number of mesodermal progenitors in the coronal suture and invasion of some NCC into the parietal osteogenic domain. In both conditional *En1*-mutants, the caudal shift of the coronal suture is just mild. In a more extreme condition, embryos carrying mutations in *Fuz*, a regulator of ciliogenesis, develop a single bone instead of frontal and parietal bones. This phenotype occurs by the caudal expansion of NC-derived frontal bone cells due to increased *Fgf8* expression in the cephalic region and a consequent reduction or absence of mesoderm-derived parietal cells. A similar although less severe phenotype was observed in mutants of the SHH effector GLI3 (Tabler et al., 2013; Tabler et al., 2016). These data indicate that the caudal shift of the coronal suture in *En1* conditional mutants or the caudal expansion of the NC observed in *Fuz*-null mutants depends on abnormal FGF and SHH signalling.

Although PLPP3 deficiency can alter the cephalic expression of *Fgf8* (this work) and potentially HH signalling (see below), it is contrasting that in our CKO-mutants the shift of the coronal suture occurs in anterior direction, suggesting that additional mechanisms for the positioning of this structure are at play. In mice, it has been suggested that positioning of the coronal suture is determined by: (1) the apposition front of tissues of different embryological origin (NCC/mesoderm); (2) inductive signals during neural tube development from the neural inductive centre zona limitans intrathalamica (ZLI) in alignment with the telencephalon/diencephalon boundary between E10.5-11.5 (Teng et al., 2019). Our data support the first hypothesis to a great extent. The increased cell death in the SOM of CKO around E10.5 reduced the NC-territory thereby allowing the anterior expansion of cells of mesodermal origin. This mesodermal population must include *En1*-expressing coronal suture progenitors, since the coronal suture mesenchyme is maintained in the CKO. These results support that signals for establishing the position of the coronal suture in the mouse arise where NC and mesodermal derivatives interact in the supraorbital region around E10.5; although additional experiments will be necessary to rule out the participation of other cues originating from the ZLI.

The altered position of mesoderm-derived tissues in CKO embryos was not restricted to the frontal-parietal boundary. At E14.5, the mesoderm-derived tectum transversum of the chondrocranium was also positioned forward, and NC-derived dermal tissue was consistently reduced in the temporal region, adjacent to the neural crest–mesoderm boundary. Thus, PLPP3 deficiency in NCC appears to widely affect the spatial organization of cranial tissues of different embryonic origins, besides the positioning of the coronal suture. Although our data suggest that these phenotypes mainly originate from the loss of NC-derived cells, other contributing mechanism should be explored, such as active mesodermal expansion or altered tissue displacement. The frontonasal phenotype is also associated with increased LysoTracker staining in the medial and lateral prominences between E10.5-11.5. This localized alteration in survival of NC-derived cells accounts for the frontonasal narrowing and hypotelorism observed in CKO.

The precise mechanism altering the survival of CKO in nasal prominences and NC/mesoderm boundaries remains to be established, however, it is likely that dysregulation of extracellular bioactive lipid signalling is involved given the expression pattern of *Plpp3,* of lipid synthesizing enzymes and of several LPA and S1P receptors in the SOM and nasal prominences around E10.5-11.5 (Fig. S5; Ji et al., 2021; Ohuchi et al., 2007; Yokoyama et al., 2009; Meng and Lee, 2009). Interestingly, *Lpar1* mutants show craniofacial dysmophology in mouse and zebra fish (Contos et al., 2000; Nishioka et al., 2016). Mutant mice show a shorter and wider snout with ocular hypertelorism, misalignment of chondrocytes and precocious ossification of the cartilage between the presphenoid and basisphenoid bones, both of neural crest origin; similar phenotypes are produced by a hypomorph/null allele of *Enpp2* (Contos et al., 2000; Nishioka et al., 2016). Moreover, 4% of *Lpar4*-deficient individuals at E18.5 show agnathia (small or absent mandible) (Sumida et al., 2010). These data highlight the relevance of LPA-mediated signalling for craniofacial development. However, given the apparently opposing phenotypes found in our study (i.e. hypotelorism vs hypertelorism), it is possible that some phenotypic features arise from overstimulation and/or desensitization of single or multiple LPA and/or S1P receptors. Alternatively, other PLPP3 activities independent of lipid receptors in the membrane could be playing an important role, namely, through the deregulation of intracellular lipids or the lack of PLPP3-integrin binding.

Midfacial hypoplasia and hypotelorism without ventral midline brain abnormalities are considered mild presentations of holoprosencephaly (HPE). HPE was initially associated with mutations in SHH and other regulators of the HH pathway, although nowadays it is established that alterations in TGFβ and FGF signalling are also common genetic causes of this congenital disorder (Rossler et al, 2018). For more than two decades, it has been known that the primary cilium is essential for HH signalling (Huangfu et al., 2003). More recently, this structure has been identified as a signalling hub for a plethora of signalling pathways among which are FGF, WNT-PCP, TGFβ/BMP and GPCR pathways (Mill et al., 2023) including LPAR1 receptors (Loskutov et al., 2018). LPA/Lpar3 mediated signalling regulates cilia number and volume in the zebra fish Küpffer’s vesicle (Lai et al., 2012) and LPA/LPAR1 signalling mediates serum induced cilium disassembly in retinal pigmentary epithelial cells and cilium length in murine neural stem cells (Hu et al., 2021). These observations evidenced the importance of regulation of LPA-mediated signalling in cilia function and potentially in the aetiology of ciliopathies. Notably, *Plpp3* (which dephosphorylates LPA), is included in a curated database of ciliary genes (Elliott et al., 2023) https://research.cchmc.org/Ciliome_Gene_Expression/) and is 1 of 13 confirmed negative regulators of cilium length identified in a screen for modulators of primary cilium formation and length (Kim et al., 2010). Together, these observations raise the possibility that loss of *Plpp3* expression in NCC may alter HPE-associated signalling in the frontonasal area. This could be achieved not only through the regulation of LPA availability, but through changes in intracellular lipids modulating the activity of HH signalling components, such as phosphatidic acid (Zhang et al., 2023) or lipids required for primary cilia formation, such as ceramide (Wang et al., 2009). Then, it would be interesting to test if altered cilium-dependent signalling contributes to the craniofacial phenotype observed in our *Plpp3* CKO. Given the potential complexity of the mechanisms originating the phenotypes described, we are currently designing multi-omics strategies (including spatial lipidomic) and *in vitro* approaches to determine how is that PLPP3-deficiency specifically produce the observed phenotypes.

Our findings raise the possibility that *PLPP3* might contribute to the craniofacial dysmorphic features observed in patients with genomic variants involving this gene. *PLPP3* maps to chromosome 1p32.2 and microdeletions involving this region are rare. Analysis of the DECIPHER GRCh38 database (DatabasE of genomiC variation and Phenotype in Humans using Ensemble Resources; Firth et al., 2009) indicated that in humans, *PLPP3* has a very high probability of loss-of-function intolerance (pLI=1.00) and low tolerance to mutations (LOEUF=0.4). Phenotypes described in some of the 18 patients with genomic variants including *PLPP3* (duplications and heterozygous deletions), suggest NCC-development involvement (protruding ear, abnormal facial shape, bifid uvula, abnormal pinna morphology, deep philtrum, low-set ears, prominent nasal bridge, wide intermaxillary distance, abnormal dentition, anteverted nares, broad forehead, cupped ear, short nose, narrow nose) (Firth et al., 2009). Genomic variants in several of these patients include genes previously associated to abnormalities in nervous system development such as *DAB1* and *NFIA*. Interestingly, deletions containing *DAB1* but not *PLPP3*, lack NCC-associated phenotypes (Firth et al., 2009) and as shown in figure 10, *DAB1* does not include craniofacial superenhancer-regions. Five clinical reports involving 1p32.2 without NFIA deletion have been described, all with craniofacial dysmorphism. Among them, one patient with a mapped interstitial 1p32.2p32.3 microdeletion (5.4 Mb) showed macrocephalia, hypertelorism, telecanthus, prominent philtrum, low set ears, cleft lip and palate, downslanting palpebral fissures, bilateral ptosis, simplified ear structures, bulbous nose, and dental crowding (Mulatinho et al., 2008). Other patient with 1p32.2p33 microdeletion (6.4 Mb) showed bilateral membranous choanal atresia, flat nasal bridge, small nose, anteverted nares, retroghnathia and microcephaly (Kehrer et al., 2015). Another case encompassing 1p32.1p32.2 (10 Mb) exhibited micrognathia, high palate, long face, frontal bossing, prominent philthrum, dental crowding, squint, patent ductus arteriosus and atrial septal defects (Jiang et al., 2020). Some of these phenotypic features are absent or not frequently observed in other 1p syndromes (micrognathia, microcephaly, dental crowding, frontal bossing, prominent philtrum, high palate, long face) (Jiang et al., 2020; Kehrer et al., 2015). *PLPP3* duplications would also alter gene dosage and therefore bioactive lipid homeostasis, hence potentially explaining the appearance of craniofacial phenotypes in some of the described patients with these genomic variants. It would be important to explore if individuals with small deletions, mapping to regions containing craniofacial enhancers (Fig. 10C; enhancer hs1622, Vista Enhancer Browser; or patients 479909 and 571027 in the DECIPHER database) show specific facial features or dysmorphisms. Finally, a Genome Wide Association Study identified 206 genomic variants influencing normal-range facial variations (White et al., 2021). Among those, rs199971562 maps to 1p32.2 within the intergenic region shown in figure 10. This variant, was associated with having a role in the morphology of a facial segment comprehending the forehead and eyes. The candidate gene associated to this variant was *PLPP3,* despite no clear association with craniofacial dysmorphology in humans or described animal models by that time. Our work constitutes the first evidence clearly associating this gene with craniofacial morphogenesis and supports its consideration as a contributor to human craniofacial variation.

### Experimental procedures

#### Animals

All procedures used in this project were carried out according to the protocol DEA141-18 and DEA196-22, approved by the Internal Committee for the Use and Care of Laboratory Animals (CICUAL) from the Institute of Cellular Physiology, UNAM. The *Wnt1::Cre2* transgenic line (*E2f1^Tg(Wnt1-cre)2Sor^*, Lewis et al., 2013) was used to drive recombination of our *Plpp3* conditional allele (*Plpp3^tm3Stw^*; Escalante-Alcalde et al., 2007). For lineage tracing experiments the double reporter line *Gt(ROSA)26Sor^tm4(ACTB-tdTomato,-EGFP)Luo^* was used (Muzumdar et al., 2007). For the analysis of MHB markers in global mutants and *Plpp3* expression analysis, embryos were obtained from heterozygous CD1.*Plpp3^tm2Stw^* inter-crosses (Escalante-Alcalde et al., 2009). Genotyping was carried out from yolk sac or tail biopsy as previously described (Escalante-Alcalde et al., 2009; Escalante-Alcalde et al., 2007). DNA from yolk sacs was obtained using a PCR compatible lysis buffer (50 mM KCl, 10 mM Tris pH 8.3, 2 mM MgCl_2_, 0.01% gelatin, 0.45% NP-40, 0.45% Tween 20 and 100 µg/ml of proteinase K).

#### Whole-mount in situ hybridization

Mouse embryos were fixed and processed as described ahead. All solutions were prepared using DEPC treated Milli Q water. Quick washes between steps were done for 5 mins using PBT. Embryos were freed from their extraembryonic membranes and fixed in 4% PFA in PBS 1X, pH7.4 at 4°C overnight then washed with PBS and dehydrated with a methanol gradient going from 25%, 50% and 75% in PBS with Tween 20 at 0.05% (PBT) and stored in 100% methanol at -20°C. Somite matched embryos were pooled by genotype and rehydrated on ice in a methanol gradient going from 75%, 50% and 25% in PBT. Embryos were then bleached for one hour in 2% H_2_O_2_ in PBT and permeabilized for 3 mins with 10 mg/ml of Proteinase K in PBT. Reaction was stopped incubating with glycine at 2 mg/ml in PBT for 10 mins. Embryos were post-fixed for 15 min with PFA 4%/glutaraldehyde 0.2% in PBT and then incubated for two hours at 70°C in hybridization solution (formamide 50%, SSC 5X pH 4.5, SDS 1%, yeast tRNA 50 µg/ml [Sigma R5636] and heparin 5 mg/µl in DEPC treated Milli Q water). Embryos were incubated overnight at 70°C with antisense or sense digoxygenin-UTP labelled RNA probes for *Otx2*, *Pax2*, *Wnt1*, *Fgf8* and *En2,* at a concentration of 500 ng/ml of hybridization solution. Sense probes were used as negative controls. Embryos were then washed three times at 70°C for 10 mins with hybridization solution, three times at 70° for 30 mins with washing buffer 1 (formamide 50%, SSC 5X pH 4.5 and SDS 1% in DEPC treated Milli Q water) and three times at 70°C for 30 mins with washing buffer 2 (formamide 50% and SSC 2X pH 4.5 in DEPC treated Milli Q water). Afterwards, embryos were given 3 quick washes at room temperature with TBST (Tris 25 mM pH 7.5, NaCl 0.234 g and KCl 0.02 g in DEPC treated Milli Q water) and then incubated 2 ½ hours in 1% Blocking Reagent in PBT (Roche 1096176). Embryos were then incubated overnight at 4°C with Anti-Digoxigenin-AP antibody (Roche 1093274) at a concentration of 1:2000 in 1% Blocking Reagent. Embryos were then washed at room temperature 5 times for 1 ½ hours with TBST and then overnight at 4°C. For developing, embryos were washed 3 times at room temperature for 10 mins with AP detection buffer (NaCl 100 mM, Tris pH 9.5 100 mM, Tween 1% and levamisole 2 mM in DEPC treated Milli Q water) and then with staining solution (5 ml AP buffer, 25 µl of NBT and 18.75 µl of BCIP) guarded from the light until the stain developed. The reaction was stopped with AP stop buffer (Tris 100 mM and EDTA 1 mM in DEPC treated Milli Q water), and then the embryos post-fixed in PFA 4% in PBS for one hour at room temperature (RT) and then dehydrated and rehydrated in a methanol series. Embryos were visualized and photographed using a Nikon camera mounted on a SMZ800 stereoscope (Nikon) or an AxioZoom stereomicroscope (Zeiss) in 30mm dishes containing a bed of agarose 1% in PBS.

#### RT-qPCR of markers in embryos’ cephalic region

Once each embryo was freed from the extraembryonic membranes, its head was cut off in an angle, above the heart in the ventral side and at the level of the rhombencephalon in the dorsal side, with the help of fine tweezers. Each cephalic portion was collected in a 500 µl Eppendorf tube containing 20 µl of RLT lysis buffer (RNeasy mini kit, Qiagen) and stored at -80°C while the rest of the embryo was used for genotyping as described above. Once the embryos were genotyped, 10 cephalic regions of each genotype (*Plpp3*^+/+^ and *Plpp3 ^lacZ/lacZ^*) were pooled and total RNA was extracted with RNeasy mini columns (Qiagen, cat. 74104). One µg of total RNA from each genotype was treated with DNaseI RNase-Free (Roche 04716728001) and reverse transcribed using AMV-RT system (Promega M900A). Each well contained 1X SYBR green (Applied biosystems 430915), 300 nM of forward and reverse primers and 25 ng of cDNA in a volume of 12.5µL. Amplification was performed in a StepOnePlus™ Real-Time PCR System. Triplicate PCR reactions underwent 1 cycle at 95 °C for 5 min, followed by 40 cycles of 95 °C for 15 s, 60 °C for 30 s in the thermal cycler and then the melt curve was carried out with standard conditions.

The following primers were used: *Fgf8* forward primer 5’-AGA AGA CGG AGA CCC CTT CG-3’, and *Fgf8* reverse primer 5’-TGA ATA CGC AGT CCT TGC CTT-3’ (Linscott and Chung, 2016); *Wnt1* forward primer 5’-TCG AGA AAT CGC CCA ACT TC-3’, and *Wnt1* reverse primer 5’-TGC AAG CTC GTC CAG CTG T-3’ (Du, et al., 2016); *Pax2* forward primer 5’-ACA TCA AAT CAG AAC AGG GGA AC-3’, and *Pax2* reverse primer 5’-CAT GTC ACGACC AGT CAC AAC-3’ (Chen, et al., 2006); for internal control *actin* forward primer 5’-TAT TGG CAA CGA GCG GTT CC-3’, and *actin* reverse primer 5’-GCA TAG AGG TCT TTA CGG ATG TC-3’. To normalize the total amount of cDNAs present in reactions from total RNA, actin cDNA was co-amplified as control. The R value (Relative expression) was calculated using the following equation: R= 2^[Ct(actin) – Ct(gen X)]^. To determine the fold change of each gene, the value obtained in wild type samples was normalized to 1, and the value in mutants was expressed as fold change vs wild type. Two pools of 10 embryos were analyzed for *Fgf8* and *Wnt1*, while only one pool was assayed for *Pax2*.

#### Scanning electron microscopy

Mouse embryos were fixed by immersion in Karnovsky’s fixative [50% Glutaraldehyde (Sigma-Aldrich), 4% Paraformaldehyde (Sigma-Aldrich), 0.2M Sodium Cacodylate buffer (Electron Microscopy Sciences) pH 7.4]. Fixation was carried out for 24 hours, followed by washes with 0.1M Sodium Cacodylate buffer. Embryos were dehydrated through a series of ethanol (EtOH) baths diluted in PBS 1X. Ascending concentrations of EtOH were used: 30%, 40%, 50%, 60%, 70%, 80%, and finally, two consecutive baths of 90% and 100%, each step lasting 30 minutes. Subsequently, samples underwent desiccation to critical point with CO_2_ (SAMDRI 780A) and coated with colloidal gold (Jeol fine coat ion sputter JFC-110). Samples were observed and photographed under a scanning electron microscope (JSM-5410LV).

#### Immunofluorescence for cell death and proliferation detection

For whole-mount preparations, embryos were collected and fixed with 4% PFA in PBS for 30 minutes at room temperature followed by 3 PBS washes of 10 min. Embryos were then permeabilized for 15 min with Triton X-100 0.5 % in PBS and then blocked in BSA 5%, 0.1% Triton-100X in PBS and incubated overnight with rocking at 4°C with phospho-Histone H3 1:100 (pHH3, Cell Signaling Technologies #9701) or active cleaved Caspase-3 1:1000 (aCas-3, Cell Signaling Technologies #9661) primary antibodies diluted in blocking solution. After 3 washes with PBS, 20 min each, secondary antibody chicken anti-rabbit Alexa Fluor 488, (Invitrogen A21441) was then diluted in blocking solution at 1:500, and incubated at room temperature for 1.5 hours. After 3 washes, nuclei were stained for 15 min using 1 mg/ml Hoechst 33342 (Molecular Probes H1399) diluted 1:1000 in PBS. Embryos were mounted in custom made chamber slides with Vectashield (Vector H1000) and observed under a confocal/two photon microscope LSM710 (Zeiss).

For immunofluorescence in sections, embryos were fixed overnight, washed with PBS, equilibrated in sucrose 30% in PBS and then, six series of coronal sections of 40 µm thick, from each embryo’s head, were obtained in a CM1900 cryomicrotome (Leica) and mounted in Superfrost® Plus slides. Each of the six series contained a representative sampling of all the levels sectioned. Slides (series) were unmasked with Citrate Buffer (Invitrogen #005000 or Vector #H-330) using a pressure cooker at 30 psi for 8 min. Unmasking treatment also denatured EGFP in embryos from lineage tracing experiments. Slides were then treated with glycine 0.1 M in PBS for 15 min at RT, 2 quick PBS washes and incubated 1 hr with 1% Blocking Reagent (Roche, #11096176001), 0.2% Triton X-100 in PBS. Primary antibodies were diluted in blocking solution and incubated 24 hrs at 4 °C followed by 3 washes 15 min each. Slides were incubated for 2 hr with secondary antibody diluted in blocking solution, washed, counterstained with Hoechst 33342 as indicated above and mounted with Fluoromount^TM^ aqueous mounting medium (Sigma-Aldrich, #F4680). For lineage tracing analysis, one series of each individual was untreated but counterstained and mounted as indicated above. Images were acquired using an LSM 800 confocal microscope (Zeiss).

#### Quantification of pHH3 positive cells in whole embryos

Matched stage *Plpp3*^+/+^, *Plpp3*^+/*lacZ*^ and *Plpp3 ^lacZ/lacZ^* embryos around E8.5-E9.0 were stained as indicated above. Dorsal views of the cephalic neural folds were acquired using a Zeiss LSM710 two photon/confocal microscope under a 20x objective. Z-planes had a thickness of 5 µm per slice. Substacks containing planes with mitotic neuroepithelial cells were selected to quantify the number of pHH3 positive cells. Images were processed using Image J. We applied a Gaussian filter with a sigma radius of two to smoothen the edges of the cells and neuroepithelium. The area of the neuroepithelium analysed was manually segmented and then quantified. The 3D Objects Counter plug-in was used to identify and quantify positive cells. Cells were segmented using a threshold and a size filter set to a minimum of 72 voxels to exclude elements smaller than a cell. The number of pHH3 positive cells counted was divided by the area of the analysed neuroepithelium and reported as cells per 2500 µm^2^ corresponding to a square of 50 x 50 µm. The data were analysed using the GraphPad Prism7 software. The Kolmogórov-Smirnov normality test was applied. Since one of the data sets was too small, a non-parametric One-way ANOVA, Kruskal Wallis, and the Dunn multiple comparison *post hoc* test were used to evaluate differences between genotypes. Histogram shows median and range plotted.

#### Quantification of pHH3 and aCas-3 positive cells in sections

Images acquired in .czi format were analyzed using Fiji (Schindelin et al., 2012). Regions of interest (ROI) were manually delineated using the polygon tool. The area of each ROI was measured, and the number of pHH3+ or active caspase-3+ cells within each region were manually quantified using the cell counter plug-in. The data were expressed as the number of marker positive cells/mm^2^. Equivalent anatomical levels were analyzed in control and CKO embryos. To avoid counting blood as false-positive cells, images were acquired in 3 channels (Alexa-488, tdTomato, Hoechst 33342), where red blood cells appeared slightly positive in both the red and green channels and were excluded from the quantification.

#### Whole mount skeletal staining

Neonate skull skeletal preparations were performed essentially as described previously (Rigueur and Lyons, 2014). Briefly, after euthanasia, heads were scalded and skin, eyes and excess tissue removed, followed by fixation in 95% EtOH overnight (ON) at RT. Tail biopsy from each individual was used for genotyping. Samples were permeabilized in acetone, stained in Alcian blue solution, destained with EtOH, pre-cleared with 1% KOH and then stained with Alizarin red solution. Samples were cleared with 50% glycerol:50% KOH (1%) and transferred to 100% glycerol for image acquisition and long-term storage.

#### Morphometric measurements

Image acquisition of skeletal preparations and embryos (bright field or fluorescence) was made under an AxioZoom V16 stereomicroscope (Zeiss) and measurements between the established landmarks were made using the ZEN2.6 program (Blue edition, Zeiss) and the Graphics line tool.

#### Histochemistry for Alkaline Phosphatase (AP)

E14.5 embryos were fixed with 4% PFA for 1 h washed 3 times with PBS and sectioned as described above to remove the brain and expose the developing frontal and parietal bones. Head halves were washed for 1 h with AP-detection buffer pH 9.5 (0.1 M NaCl; 0.1 M Tris-base; 1% Tween-20; in ddH_2_O). After a second wash of 15 min, samples were incubated in AP detection solution (5 ml AP-detection buffer; 1 Fast Red TR tablet [ROCHE, 1496549]; 20 mg of naphthol AS-TR phosphate disodium [SIGMA, N6125-1G]) at 37 °C covered from light with gentle shaking. To stop the reaction, samples were washed 3 times for 30 min with PBS, post-fixed with 4% PFA overnight at 4°C and washed 3 times with PBS.

#### β-galactosidase staining of whole embryos and embryo’s heads

Embryos were stained essentially as previously described (Escalante-Alcalde et al., 2009). Briefly, embryos between E9.5-11.5 were immersed in fixative solution (0.2% glutaraldehyde; 5 mM EGTA, pH 8; 2 mM MgCl_2_; 0.1 M phosphate buffer, pH 7.3), washed 3 times for 30 min with detergent rinse (2 mM MgCl_2_; 0.01% sodium deoxycholate; 0.02% IGEPAL; 0.1 M phosphate buffer, pH 7.3) and then incubated in staining solution at 37°C (2 mM MgCl_2_; 0.02% IGEPAL; 0.01% sodium deoxycholate; 5 mM K_4_Fe(CN)_6_; 5 mM K_3_Fe(CN)_6_; 0.1 M phosphate buffer, pH 7.3; 1 mg/ml X-gal; 20mM Tris-HCl, pH 7.5) until the staining developed. For staining of E14.5 heads, embryos were first immersed in fresh fixative solution for at least 40-50 min at room temperature. Embryos were then sectioned below the forelimbs and fixed for an additional 15 min. Then, a midsagittal cut was made through each head and both halves were washed three times for 30 min with detergent rinse solution. In the meantime, the brain was removed to expose the area of the developing frontal and parietal bones. The samples were then incubated at 37°C in staining solution. Finally, samples were washed 3 times with 1X PBS, 30 min each. If crystals were formed, samples were previously washed with 70% ethanol before image acquisition.

#### Detection of cell death in whole embryos by Lysotracker staining

Indirect detection of cell death was essentially made as previously described (Fogel et al., 2012), with some modifications. Briefly, E10.5-11.5 embryos were dissected and individually collected in a 24-well plate in Leibovitz L-15 medium on ice. Embryos were incubated in 0.5 ml of L-15 containing Lysotracker^TM^ Red DND-99 0.2 µM to increase the signal-to-noise ratio. Embryos were incubated for 45 min at 37 °C with regular mild shaking then washed with PBS 4 times, fixed overnight in 4% PFA, dehydrated through methanol series and stored at -20 °C protected from light as indicated the protocol. Embryos were cleared and individually observed in glycerol 70% in PBS, in a deep concave-depression glass plate under an AxioZoom stereomicroscope (Zeiss) outfitted with Lysotracker Red and EGFP filters using the same exposure conditions for each batch of embryos.

#### Synteny, epigenomic map and embryo scRNA molecular maps

Mouse-human genome synteny map was performed using the JAX Synteny Browser (Kolishovski et al., 2019). Visualization of the human craniofacial tissue epigenomic data and chromatin state segmentations (GEO: GSE97752; (Wilderman et al., 2018) was performed loading the Cotney lab human craniofacial hub on the UCSC Genome Browser (Casper et al., 2026). Expression of the genes of interest in the single-cell molecular map from E8.5 embryos were produced in the interactive server https://crukci.shinyapps.io/mousegastrulation2018/ (Pijuan-Sala et al., 2019). Plots were UMAP projections plotted by cell type.

#### NCC migration assay

Neural plate border regions were explanted and cultured essentially as described previously (Gonzalez Malagon et al., 2019). After 24 hours of culture, explants were fixed and NCC identified by SOX10 immunofluorescence and images acquired under a LSM800 confocal microscope (Zeiss). NCC migration distance was measured between the neural tissue border and the farther migration front of NCC using the ZEN2.6 program (Blue edition, Zeiss) and the Graphics line tool.

## Acknowledgements

We would like to thank Yazmín Ramiro Cortés and Gerardo Rodrigo Perera for their help and support on the two-photon image acquisition of immunofluorescences. We would also like to thank the Microscopy, Molecular Biology, Histology and Computer Units for lending equipment. Special thanks to our Institute’s animal facility staff for careful maintenance of our mouse colony. To Silvia Espinosa Matías from the Faculty of Sciences UNAM, for her support in the acquisition of SEM images; Lourdes Massieu Trigo for providing us with Lysotracker Red. This work was supported by grants from: DGAPA-UNAM-PAPIIT IN225723 and IN225226; SECIHTI CB-2015-01-250613 and CBF-2025-I-777 to D.E.A.

---

**Supplemental figure S1. Single cell molecular map of E8.5 embryos.** Expression of the indicated genes in midbrain/hindbrain (red oval) and neural crest cells (purple area) (Pijuan-Sala *et al,* 2019; https://marionilab.cruk.cam.ac.uk/MouseGastrulation2018/).

**Supplemental figure S2. NCC migration assay.** Explants of neural tubes (nt) from wild-type (**A**) and homozygous mutant (**B**) E9.5 embryos, cultured 24 hrs. Ten measurements per explant (n=3) were made of the NCC-outgrow migration from the neural tube border (dashed grey lines). NCC were identified by nuclear immunostaining of SOX10 (green). Scale bar= 50 μm.

**Supplemental figure S3. Morphometric measurements of neonate skulls.** A) localization of the landmarks (Table S1) used for morphometric analyses. B) No significant difference was observed in the nasal bone’s length between genotypes. In CKO skulls, a consistent reduction of the distance between the measured points was registered in the frontal bone (C-G), whilst in the parietal bone was increased (H). (I) The distance between points IP and IP’ is indicative of the loss of tissue of neural crest origin in the midanterior part of the interparietal bone. *** and **** corresponds to P= 0.0001. Dunnett’s one-way ANOVA.

**Supplemental figure S4. Morphometric analysis of E11.5 and E14.5 embryos.** A) Measurements were made as indicated in the scheme, using the ZEN software (Zeiss). B) Representative images and registered measurements in (C). Differences in E11.5 CKO embryos are detectable (**P0.0012, *P0.012 and 0.02) and become more pronounced in E14.5 (****P<0.0001). An unpaired Student’s *t* test was applied. E11.5: *F/+;Cre* n=10 or 7; *F/F;Cre* n=6. E14.5: *F/+;Cre* n=9; *F/F;Cre* n=13. Scale bars: E11.5=500 µm; E14.5 lateral view=2000 µm; E14.5 frontal view=1000 µm. Asterisks in E14.5 embryos indicate the ear/pinna; arrows point to abnormalities (changes in eyes direction, loss of frontonasal tissue and increased vascularization in the frontal bone area).

**Supplemental figure S5. Expression patterns.** Expression of *Plpp3*-(Escalante-Alcalde et al 2009), *Lpar1*-, *Lpar4-* and *Lpar*-β-galactosidase reporter alleles (from the IMPC, Wilson et al, 2026) and in situ hybridization of *Enpp2* (EMBRYS 18606, Yokoyama et al., 2009) at the indicated gestation days. E14.5 embryos sectioned in half were double stained for β-galactosidase and alkaline phosphatase, lateral and midsagittal views. fr, frontal; p, parietal; white arrowhead, tectum transversum; AP, alkaline phosphatase (red color).

